# Depolarization block in substantia nigra pars reticulata neurons drives dystonia in a mouse model

**DOI:** 10.64898/2026.09.15.751818

**Authors:** Olivia K. Barnhill, Kasandra Romo, Alexandra B. Nelson

## Abstract

Dystonia is a common movement disorder characterized by involuntary co-contraction of antagonist muscles, resulting in abnormal postures. The brain often appears grossly normal, suggesting dysfunctional neural activity as a cause. The basal ganglia are implicated in dystonia, but the causal patterns of neural activity remain unknown. We used a transgenic mouse model of human paroxysmal nonkinesigenic dyskinesia (PNKD), in which ethanol triggers dystonic attacks, to test the role of the substantia nigra pars reticulata (SNr), the primary output nucleus of the basal ganglia in rodents. The firing of SNr neurons was profoundly reduced *in vivo* during dystonic attacks. Multiple physiological approaches indicated that in PNKD mice, ethanol leads to depolarization block in SNr neurons. Indeed, pulsatile optical activation of inhibitory striatal inputs to SNr rescued firing and attenuated dystonia. In line with theories of basal ganglia dysfunction, these results suggest reduced activity in basal ganglia output nuclei causes dystonia.

## Main

Dystonia is a common movement disorder characterized by involuntary co-contraction of antagonistic muscles, producing abnormal movements and postures^1^. Dystonia often interferes with normal movement, leading to disability. While botulinum toxin is helpful for focal dystonias (affecting one muscle group), pharmacological therapies for generalized dystonia (affecting many muscle groups) are often ineffective. Our lack of mechanistic understanding of dystonia limits therapeutic development.

In most patients with dystonia, the brain appears structurally normal, suggesting aberrant neural activity may underlie the disorder. Multiple neural circuits have been implicated^2,3^, but dystonia is classically considered a disorder of the basal ganglia^4–7^. In dystonia associated with brain lesions, the basal ganglia are most frequently involved^8,9^. Furthermore, basal ganglia deep brain stimulation (DBS) is therapeutic for many patients with dystonia^10–13^. While these clinical observations suggest that the basal ganglia play a critical role in dystonia, the specific changes in neuronal activity that cause the disorder remain unknown.

The two main output nuclei of the basal ganglia, the globus pallidus pars interna (GPi) and the substantia nigra pars reticulata (SNr), contain GABAergic projection neurons with high tonic firing rates. A longstanding model of basal ganglia function posits that during voluntary movement, specific ensembles of GPi/SNr neurons pause firing to release specific movements, while others actively inhibit alternative motor programs^14–18^. Reduced or abnormally patterned GPi/SNr firing could disinhibit competing motor programs, leading to dystonia^6,7,9,19–21^.

Testing the role of basal ganglia dysfunction in dystonia has been difficult due to limitations in human neural recordings and animal models. For safety reasons, implantation of DBS devices in people with dystonia is often performed under general anesthesia, which alters neural firing properties and prevents moment-to-moment correlation of dystonia with neural activity. Available intraoperative recordings reveal lower firing rates in basal ganglia output nuclei compared to healthy non-human primates or people with Parkinson’s disease^22–26^, but establishing a causal role is difficult in human subjects. Rodent models could fill this gap. To date, such models fall into two categories: “etiologic models”, which reproduce a known human cause such as a gene mutation (e.g. DYT1)^27–33^, or “phenotypic models”, which display a motor syndrome resembling dystonia^34–39^. Unfortunately, few if any rodent models of dystonia combine etiologic and phenotypic validity, making it challenging to explore the underlying causes of dystonia.

Here, we take advantage of a transgenic mouse model of a human disorder, paroxysmal nonkinesigenic dyskinesia (PNKD)^40–42^. PNKD is a rare autosomal dominant disorder caused by mutations in the *PNKD* gene^40,43–45^, characterized by attacks of involuntary movements, including dystonia, in response to triggers such as alcohol, caffeine, or severe stress^46,47^. Importantly, mice harboring human *PNKD* mutations exhibit a similar phenotype: attacks of involuntary movements triggered by alcohol, caffeine, or severe stress^41^. Alcohol tends to trigger more severe dystonic attacks^41^. With both etiologic and phenotypic validity, PNKD mice offer an opportunity to identify the physiological substrates of dystonia. In PNKD mice, we found a profound reduction in SNr firing that closely mirrored the time course of ethanol-induced dystonia. Both *in vivo* and *ex vivo* physiology indicate reduced SNr activity is mediated by depolarization block, where neurons depolarize so dramatically that sodium channels supporting action potential electrogenesis are largely in a state of voltage-dependent inactivation. Finally, we provide evidence that this loss of SNr activity causes dystonia: pulsatile activation of inhibitory inputs to the SNr led to recovery of SNr spiking and attenuated dystonia. Our data demonstrate the critical role of the basal ganglia and suggest an underlying mechanism for the loss of SNr firing during PNKD dystonia.

## Results

### SNr firing rates are profoundly reduced during dystonia

To examine the physiological substrates of dystonia, we used PNKD mice, which display normal movements at baseline but develop dystonia in response to ethanol^41^. We administered intraperitoneal (i.p.) ethanol (1.5 g/kg) to PNKD mice and wild-type (WT) littermates. A blinded rater scored dystonia using a time-based 0–4 severity scale on hindlimbs, forelimbs, tail, and trunk, summed for a composite dystonia score of 0–16 (Fig. 1a,b; see Figure Legends and Table S1 for all statistical details). WT controls did not develop dystonia before or after ethanol injection (Fig. 1c). In contrast, PNKD mice developed pronounced dystonia peaking 10–70 minutes after ethanol injection (Fig. 1d; Supplementary Video 1).

**Fig. 1:**
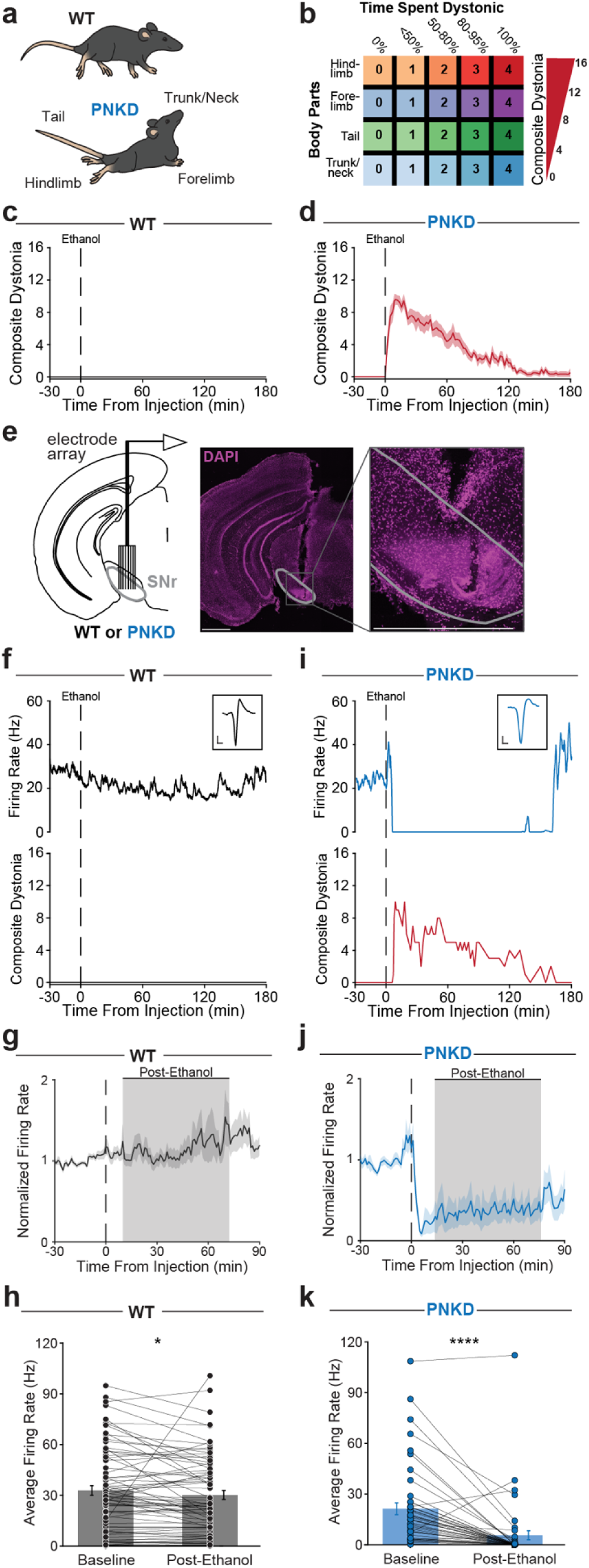
SNr firing rates are profoundly reduced during dystonia. **a,** Body parts that were observed for signs of dystonia in both wild-type (WT) and paroxysmal nonkinesigenic dyskinesia (PNKD) mice. **b,** Composite dystonia scoring system. **c,d,** Observed dystonia before and after i.p. injection of 1.5 g/kg ethanol in WT (c) and PNKD (d) mice (WT: N = 12, PNKD: N = 12, Mann-Whitney U test comparing genotypes, \*\*\*\**P* < 0.0001). **e,** Schematic of electrode recording configuration in the SNr (left) and representative postmortem tissue showing recording sites (middle) with the boxed region magnified to show detail (right). White scale bars represent 1 mm. **f,** Representative SNr neuron firing rate (top) aligned to dystonia severity (bottom) before and after ethanol injection in a WT mouse. Inset contains average waveform from this representative neuron. Scale bar represents 100 µV and 500 µS. **g,** Average normalized firing rate of SNr neurons in WT mice before and after ethanol injection (N = 7 mice, n = 75 neurons). **h,** Comparison of WT SNr neuron firing rates during baseline (0–30 minutes prior to ethanol injection) and post-ethanol (10–70 minutes after ethanol injection) periods (N = 7, n = 75, Wilcoxon signed-rank test, \**P* = 0.0395). **i,** Representative SNr neuron firing rate (top) aligned to dystonia severity (bottom) before and after ethanol injection in a PNKD mouse. Inset contains average waveform from this representative neuron. Scale bar represents 100 µV and 500 µS. **j,** Average normalized firing rate of SNr neurons in PNKD mice before and after ethanol injection (N = 6, n = 46). **k,** Comparison of PNKD SNr neuron firing rates during baseline and post-ethanol periods (N = 6, n = 46, Wilcoxon signed-rank test, \*\*\*\**P* < 0.0001). Data shown as mean ± SEM (c,d,g,h,j,k).

Clinical observations and longstanding models of basal ganglia circuit function make the activity of basal ganglia output neurons a strong candidate in mediating dystonia^4–8,10,23,48^. We investigated the neural correlates of dystonia in the primary rodent basal ganglia output nucleus, the substantia nigra pars reticulata (SNr). We recorded SNr single-unit activity before and after ethanol injection in freely-moving PNKD mice and WT littermates (Fig. 1e–k; Fig. S1). Consistent with prior studies^49–52^, SNr neurons in both genotypes showed narrow waveforms and high tonic firing rates during the baseline period (Fig. 1f–k). Average baseline SNr firing rates were modestly lower in PNKD versus WT mice (Fig. 1h,k), indicating that even during normal movement, spiking in SNr neurons is altered in PNKD mice.

We next tested how ethanol changed SNr firing. In WT mice, ethanol led to slightly lower SNr firing rates (Fig. 1f–h). However, in PNKD mice, ethanol profoundly reduced SNr firing (Fig. 1i– k). The time at which SNr firing fell below the baseline rate correlated with the onset of dystonic movement (Fig. S2a and see example Fig. 1i). Conversely, in extended recordings where dystonia resolved completely, SNr firing also recovered (Fig. 1i; Fig. S2b). These findings indicate that dystonia correlates with a profound reduction in SNr firing in PNKD mice.

### PV-SNr firing is profoundly reduced during dystonia

The SNr contains several cell types, the most numerous of which express parvalbumin (PV-SNr)^50,53^. These GABAergic projection neurons fire at high tonic rates^49^ and inhibit downstream motor areas^50,54^. We next tested whether the firing of PV-SNr neurons also fell during dystonic attacks. To identify PV-SNr neurons, WT;PV-Cre or PNKD;PV-Cre mice were injected with an AAV encoding Cre-dependent channelrhodopsin-2 (ChR2)-eYFP and implanted with an optrode in the SNr (Fig. 2a). We recorded single units before and after ethanol injection, opto-tagging PV-SNr neurons at the end of each session (after motor recovery). PV-SNr cells showed consistent, short-latency spiking in response to blue light pulses (Fig. 2b). We first compared the baseline firing rates of PV-SNr neurons in WT and PNKD littermates. As expected, PV-SNr neurons had high baseline firing rates in WT animals (Fig. 2c,d). In PNKD mice, these firing rates were substantially lower (Fig. 2e,f). We next tested how the firing of PV-SNr neurons changed after ethanol administration. In WT mice, PV-SNr firing did not change during the post-ethanol period, whereas in PNKD mice, these neurons showed a dramatic reduction in firing, essentially to zero (Fig. 2c–f). These results indicate PV-SNr neurons become nearly silent during dystonic attacks in PNKD mice.

**Fig 2:**
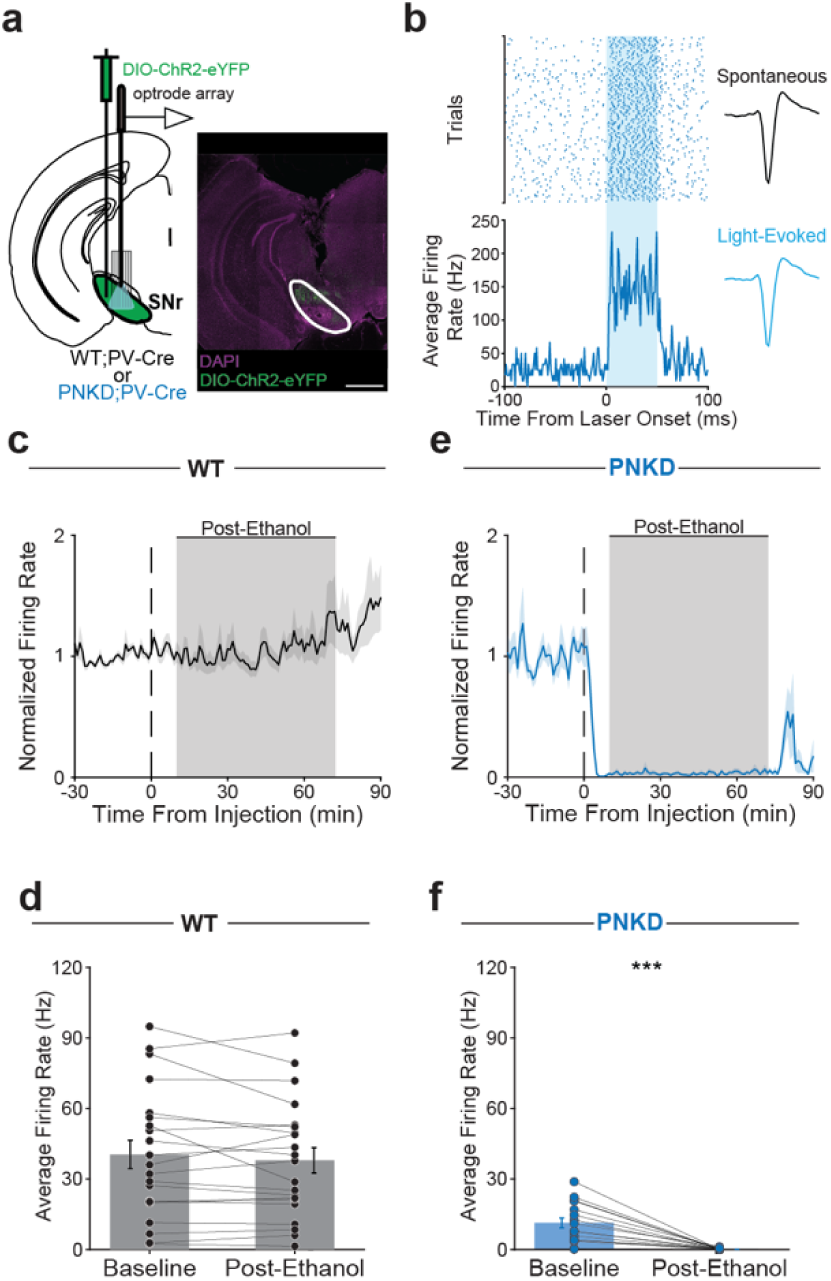
PV-SNr firing is profoundly reduced during dystonia. **a,** Schematic showing optrode recording configuration and injection of AAV encoding DIO-ChR2-eYFP in the SNr of either WT;PV-Cre or PNKD;PV-Cre mice (left). Representative postmortem coronal section showing ChR2-eYFP viral expression and optrode placement in the SNr (right). White scale bar represents 1 mm. **b,** Representative optogenetically-labeled unit, showing short-latency spiking in response to blue light across 50 trials (left) and matching spontaneous and light-evoked waveforms (right). **c,** Average normalized firing rate of SNr-PV neurons in WT mice before and after ethanol injection (N = 3, n = 21). **d,** Comparison of WT SNr-PV neuron firing rate during baseline and post-ethanol periods (N = 3, n = 21, Wilcoxon signed-rank test, *P* = 0.6639). **e,** Average normalized firing rate of SNr-PV neurons in PNKD mice before and after ethanol injection (N = 4, n = 16). **f,** Comparison of PNKD SNr-PV neuron firing rates during baseline and post-ethanol periods (N = 4, n = 16, Wilcoxon signed-rank test, \*\*\**P* = 0.0004). Data shown as mean ± SEM (c–f).

### *In vivo* signatures of depolarization block in SNr neurons during dystonia

The near-complete silencing of PNKD SNr neurons resembled depolarization block, a state in which sustained membrane depolarization prevents recovery of sodium channels from steady-state inactivation, preventing further spiking^55–59^. As sodium channel inactivation accumulates, action potentials characteristically decrease in amplitude and increase in width before the cessation of firing. We looked for hallmarks of this phenomenon in SNr single unit waveforms (Fig. 1). We compared waveforms in the baseline period and after ethanol injection but immediately preceding a significant reduction in firing rate (“Pre-Rate Decrease”; Fig. 3a–d; see Methods). We hypothesized that in PNKD recordings, SNr waveforms would show a reduction in amplitude and increase in width as compared to baseline. Indeed, PNKD SNr unit waveforms became smaller and broader over this short period (Fig. 3b–d). Although large waveform changes can affect spike-sorting, these same units resumed firing as dystonia resolved, arguing against irreversible loss of unit isolation (Fig. S2b). To confirm these changes were specific to PNKD mice, and not a general consequence of ethanol administration or our waveform selection procedure, we performed the same analysis on WT SNr neurons, sampling a matched time window (Fig. 3e,f). As expected, neither spike amplitude nor width changed in WT SNr waveforms (Fig. 3f–h). These findings are consistent with the hypothesis that in PNKD mice, ethanol causes SNr neurons to enter depolarization block, resulting in little to no firing during dystonic attacks.

**Fig. 3:**
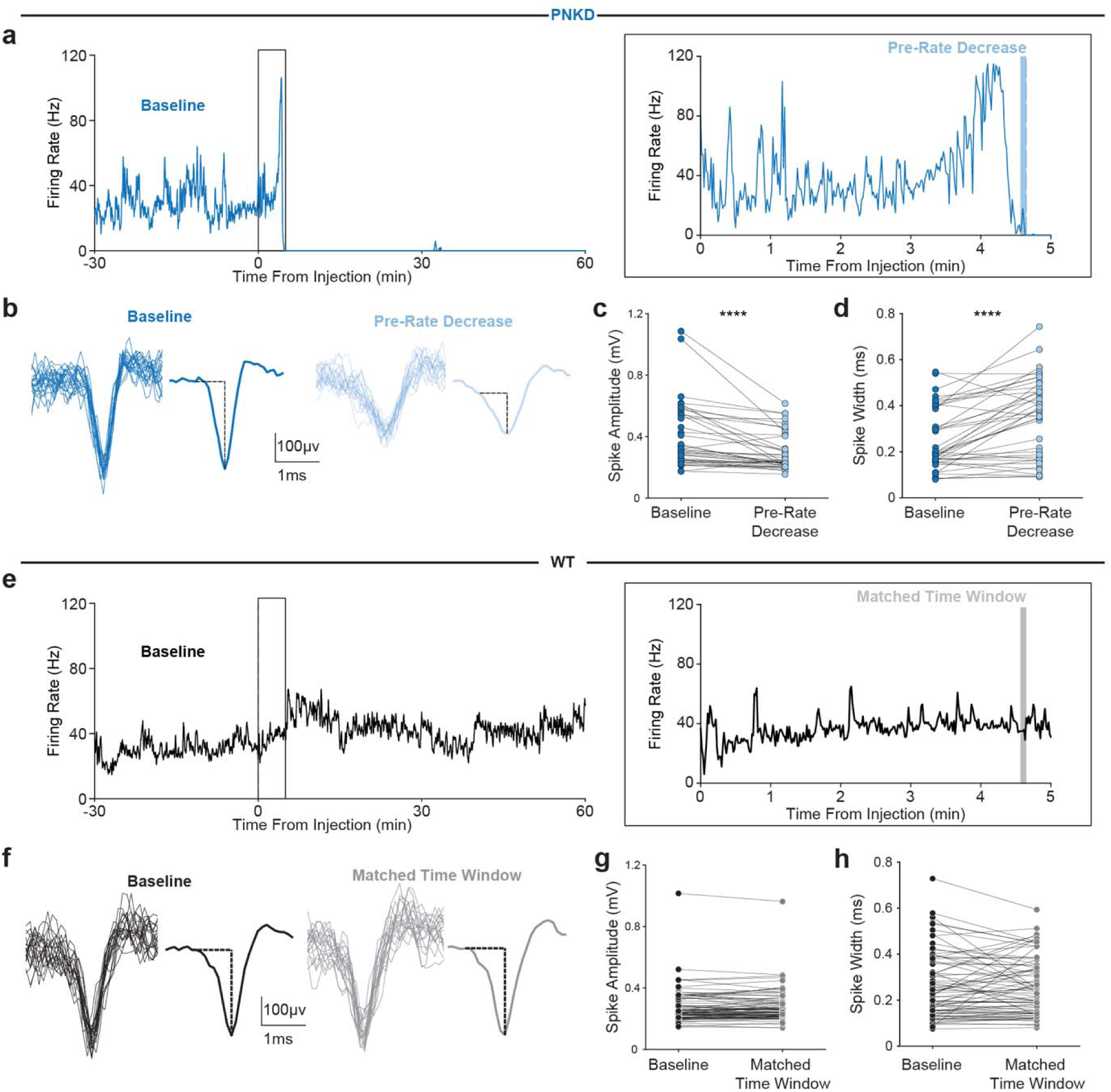
*In vivo* signatures of depolarization block in SNr neurons during dystonia. **a,** Representative SNr neuron firing rate before and after the administration of ethanol in a PNKD mouse. An expanded view of the recording 0–5 minutes after ethanol injection is shown to the right. The pre-rate decrease period is defined as the 5-second window immediately preceding either (i) the point at which firing rate dropped below the 99% confidence interval of baseline firing, or (ii) the onset of a period of ≥ 5 seconds without an action potential. **b,** 20 waveforms (individual: left, averaged: right) were randomly selected from the baseline period (10–5 minutes prior to ethanol injection, dark blue waveforms) and pre-rate decrease period shown in (a) (light blue waveforms). Waveforms shown are from a representative PNKD SNr neuron. **c,** Comparison of PNKD SNr neuron waveform amplitudes in the baseline and pre-rate decrease periods (N = 6, n = 40, Wilcoxon signed-rank test, \*\*\*\**P* < 0.0001). The spike onset was defined as the point preceding the maximal negative slope. Amplitudes were calculated from the averaged waveforms as the voltage difference between the spike onset and the negative trough. **d,** Comparison of PNKD SNr neuron waveform widths in the baseline and pre-rate decrease periods (N = 6, n = 40, Wilcoxon signed-rank test, \*\*\*\**P* < 0.0001). Widths were calculated from the averaged waveforms as the time from the spike onset to the negative trough. **e,** Representative SNr neuron firing rate before and after the administration of ethanol in a WT mouse. An expanded view of the recording 0–5 minutes after ethanol injection is shown to the right. For each neuron, a unique 5-second matched time window was randomly selected from within the aggregate window during which PNKD neurons exhibited a firing rate decrease (2–6 minutes after ethanol injection). **f,** 20 waveforms (individual: left, averaged: right) were randomly selected from the baseline period (black waveforms) and a matched time window shown in (e) (gray waveforms). Waveforms shown are from a representative WT SNr neuron. **g,** Comparison of WT SNr neuron waveform amplitudes in the baseline and matched time window periods (N = 7, n = 75, Wilcoxon signed-rank test, *P* = 0.3195). **h,** Comparison of WT SNr neuron waveform widths in the baseline and matched time window periods (N = 7, n = 75, Wilcoxon signed-rank test, *P* = 0.7453).

### SNr neuron intracellular calcium increases during dystonia

Extracellular recordings described above revealed changes in firing rate and action potential waveforms suggestive of depolarization block. To more directly assess for membrane depolarization consistent with depolarization block, we examined SNr neuronal calcium dynamics relative to dystonic attacks. High-voltage activated calcium channels, including Ca_V_2.2 channels localized to SNr dendritic domains, open at depolarized membrane potentials, and do not inactivate completely in response to prolonged depolarization^51,60^. Thus, calcium dynamics may serve as a proxy for voltage in a neuronal population experiencing depolarization. To measure intracellular calcium, we used GCaMP6s fiber photometry. The SNr of WT;PV-Cre and PNKD;PV-Cre mice was injected with an AAV encoding Cre-dependent GCaMP6s and implanted with an optical fiber (Fig. 4a,b). We then recorded bulk GCaMP6s fluorescence before and after ethanol injection. In WT mice, GCaMP6s signal declined modestly in the post-ethanol period (Fig. 4c,d), mirroring the small decrease in firing rate of SNr neurons during this period (Fig. 1h). By contrast, we saw a substantial *increase* in SNr GCaMP6s signal in PNKD mice in the post-ethanol period (Fig. 4e,f), that paralleled the decrease in SNr firing seen during dystonia (Fig. 1i). These results are consistent with the hypothesis that SNr neurons are subject to depolarization block during PNKD dystonia.

**Fig. 4:**
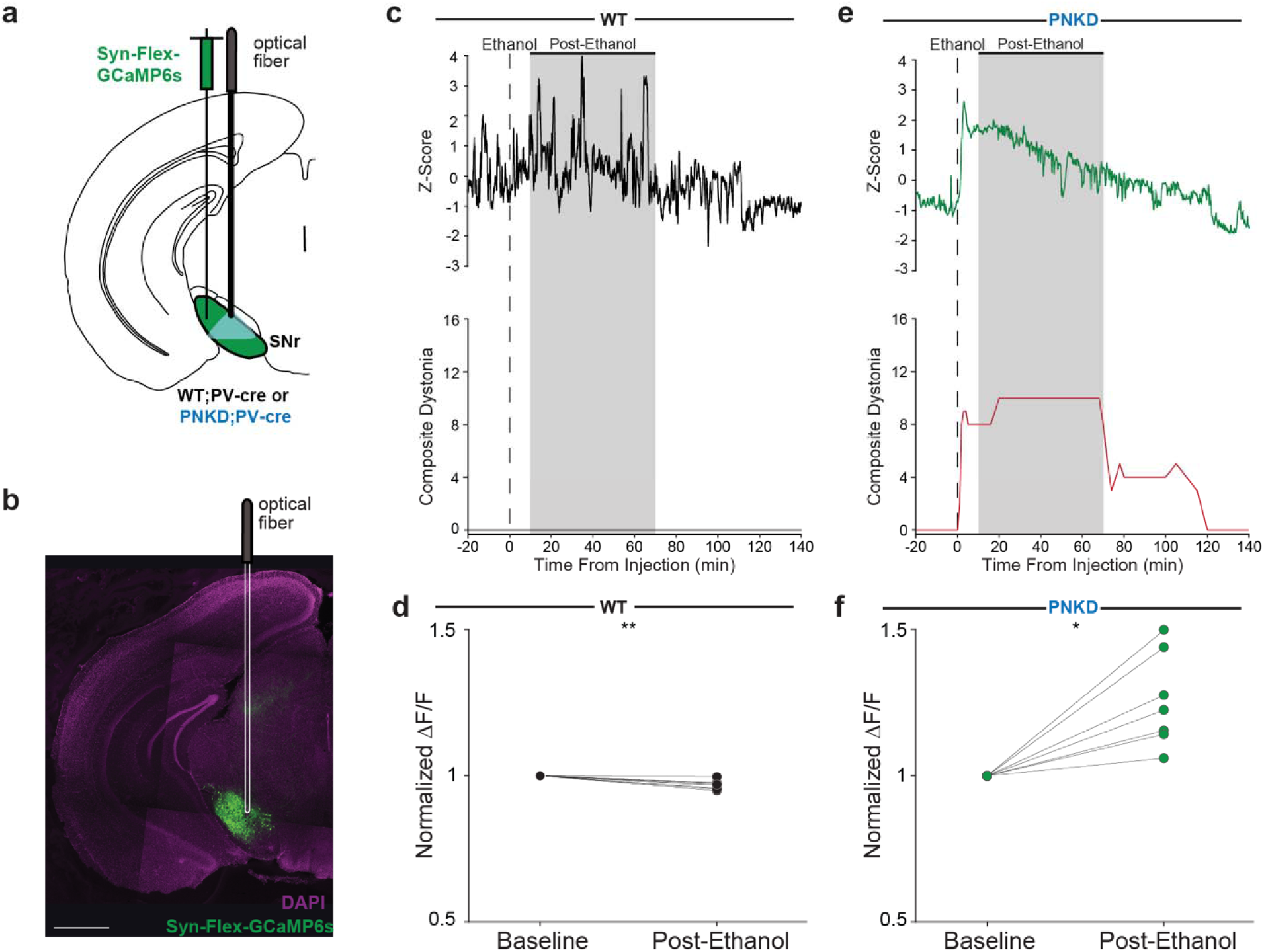
SNr neuron intracellular calcium increases during dystonia. **a,** Schematic showing injection of an AAV encoding Syn-Flex-GCaMP6s in the SNr and implantation of an optical fiber above the SNr of WT;PV-Cre or PNKD;PV-Cre mice. **b,** Representative coronal section showing the viral expression of GCaMP6s and the optical fiber placement. White scale bar represents 1 mm. **c,** Representative GCaMP6s signal, z-scored to the entire recording session (top) aligned to dystonia severity (bottom) before and after ethanol administration in a WT mouse. **d,** Comparison of WT SNr ΔF/F GCaMP6s signal, averaged within baseline and post-ethanol periods and normalized to each animal’s baseline signal (N = 9, Wilcoxon signed-rank test, \*\**P* = 0.004). **e,** Representative GCaMP6s signal, z-scored to the entire recording session (top) aligned to dystonia severity (bottom) before and after ethanol administration in a PNKD mouse. **f,** Comparison of PNKD SNr ΔF/F GCaMP6s signal, averaged within baseline and post-ethanol periods and normalized to each animal’s baseline signal (N = 7, Wilcoxon signed-rank test, \**P* = 0.0156).

### Optogenetic stimulation of PV-SNr neurons during dystonia does not increase firing

If PV-SNr neurons are in depolarization block, they should be unable to spike in response to depolarizing inputs. To test this idea, we injected AAV encoding Cre-dependent ChR2-eYFP and implanted an optrode in the SNr of PNKD;PV-Cre mice (Fig. 5a). In each mouse, we provided optical stimulation at two timepoints during the recording: once during the dystonic attack and again after dystonia resolved (Fig. 5b). During the recovery period, PV-SNr cells showed short-latency firing in response to blue light pulses (Fig. 5c,d), confirming their PV+ identity and capacity of firing in response to optical stimulation. We then examined the response of all such PV-SNr neurons to blue light pulses during the dystonic attack. These cells showed very low firing rates, and strikingly, blue light pulses were unable to elicit firing during dystonic attacks (Fig. 5e,f). These data indicate that during dystonic attacks, SNr neurons enter a state during which direct optical stimulation cannot drive firing.

**Fig. 5:**
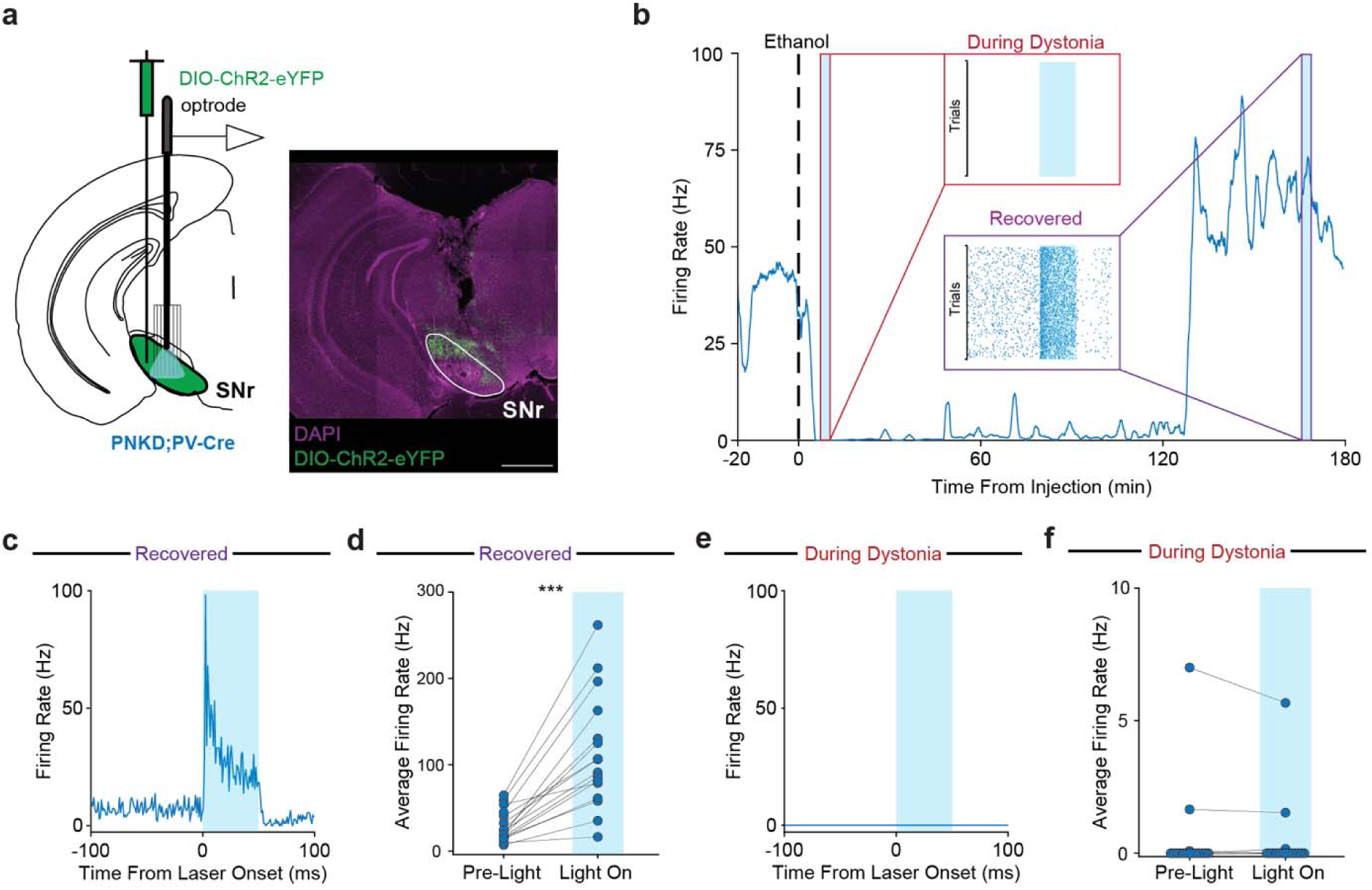
Optogenetic stimulation of PV-SNr neurons during dystonia does not increase firing. **a,** Schematic showing optrode recording configuration and injection of an AAV encoding DIO-ChR2-eYFP in the SNr of PNKD;PV-Cre mice (left). Representative postmortem coronal section showing ChR2-eYFP viral expression and optrode placement in the SNr (right). White scale bar represents 1 mm. **b,** Representative firing rate of a PNKD SNr-PV neuron that was optically stimulated during ethanol-induced dystonia and after the mouse had behaviorally recovered from the dystonic attack. An expanded view of the neuron’s response to blue light during dystonia is shown in the red box labeled “During Dystonia”. An expanded view of the same neuron’s response to optical stimulation after the mouse had recovered is shown in the purple box labeled “Recovered”. **c,** Peristimulus time histogram (PSTH) showing average firing rate during optical stimulation for the cell represented in (b) during behavioral recovery. **d,** After data collection, PV+ cells were optically identified by light-mediated increases in firing rate during the recovery period. Comparison of average firing rates during the pre-light and light-on periods during behavioral recovery (N = 5, n = 16, Wilcoxon signed-rank test, \*\*\**P* = 0.0004). **e,** PSTH showing responses from the same cell in (b) and (c) during dystonia. **f,** The same cells, confirmed as PV+ in (d), were also stimulated with blue light during dystonic attacks. Comparison of average firing rates during pre-light and light-on periods during dystonia (N = 5, n = 16, Wilcoxon signed-rank test, *P* = 0.625).

### PNKD SNr neurons have altered intrinsic properties

To investigate why PNKD PV-SNr neurons are susceptible to depolarization block, we performed cell-attached and whole-cell recordings from mCherry-positive neurons in acute SNr slices from PNKD;PV-Cre or WT;PV-Cre animals injected with AAV encoding Cre-dependent mCherry (Fig. 6a). As compared to WT mice, PV-SNr neurons in PNKD mice had lower spontaneous firing rates in both cell-attached and whole-cell modes (Fig. 6b–c). Consistent with this observation, the average membrane potential of PNKD PV-SNr neurons was slightly more hyperpolarized compared to WT (Table S2). The input resistance did not differ between SNr neurons recorded from WT and PNKD mice (Table S2).

**Fig. 6:**
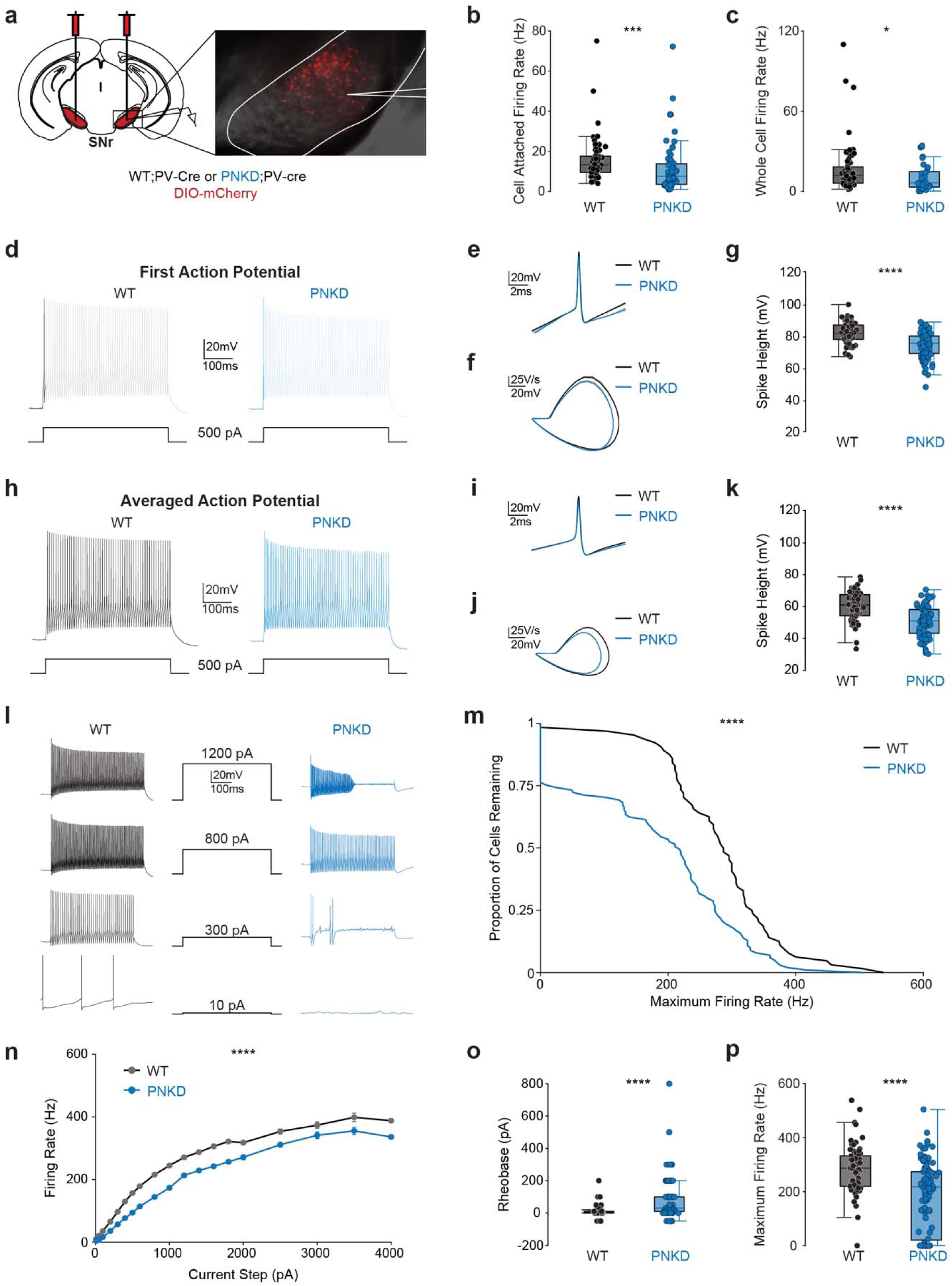
PNKD SNr neurons have altered intrinsic properties. **a,** Schematic showing bilateral injection of an AAV encoding DIO-mCherry in the SNr of WT;PV-Cre or PNKD;PV-Cre mice (left) and representative DIC image showing the patch clamp electrophysiology recording configuration. **b,** Comparison of cell-attached firing rate of WT (N = 6, n = 58) and PNKD (N = 7, n = 63) SNr neurons (Mann-Whitney U test, \*\*\**P* = 0.0002). **c,** Comparison of whole cell firing rates of WT (N = 6, n = 62) and PNKD (N = 7, n = 32) SNr neurons (Mann-Whitney U test, \**P* = 0.0188). **d–k,** Action potential shapes were analyzed from SNr neurons held at -70 mV and evoked with a 500 ms, 500 pA current step. The same recordings were analyzed two ways: using the first evoked action potential (d–g) or the average of all action potentials evoked across the sweep (h–k). **d,** Representative WT (left) and PNKD (right) responses to the current step, with the first evoked action potential used for analysis in (e–g). **e,** Group-average shape of the first action potential for WT (black) and PNKD (blue) SNr neurons. **f,** Average phase-plane plot for the first action potentials for WT and PNKD SNr neurons. **g,** Comparison of first action potential spike height in WT and PNKD SNr neurons (WT: N = 6, n = 69; PNKD: N = 7, n = 96; Mann-Whitney U test, \*\*\*\**P* < 0.0001). **h,** Representative WT (left) and PNKD (right) responses to the current step, with all evoked action potentials averaged for analysis in (i–k). **i,** Group-average of the averaged-waveform shape for WT and PNKD SNr neurons. **j,** Average phase-plane plot of the averaged waveform shape for WT and PNKD SNr neurons. **k,** Comparison of averaged-waveform spike height in WT and PNKD SNr neurons (WT: N = 6, n = 69; PNKD: N = 7, n = 96; Mann-Whitney U test*, ****P* < 0.0001). **l–p**, Summary of the input-output characteristics of WT and PNKD SNr neurons. **l,** Representative outputs from a WT (left, black) and a PNKD (right, blue) SNr neuron in response to current injections (10, 300, 800, and 1200 pA steps). **m,** Survivor curve showing the proportion of WT and PNKD SNr neurons reaching each maximum firing rate (WT: N = 6, n = 64; PNKD: N = 7, n = 101; two-sample Kolmogorov-Smirnov test, \*\*\*\**P* < 0.0001). Cells that were unable to fire action potentials across the entire 450 ms sweep at any current injection have a maximum firing rate of 0 Hz. **n,** Input-output curve showing the average firing rate responses from WT and PNKD SNr neurons in response to current injections (WT: N = 6, n = 63; PNKD: N = 7, n = 77; linear mixed-effects model, \*\*\*\**P* < 0.0001). **o,** Comparison of WT and PNKD SNr neuron rheobases from the input-output curve (WT: N = 6, n = 62; PNKD: N = 7, n = 77; Mann-Whitney U test, \*\*\*\**P* < 0.0001). **p,** Comparison of WT and PNKD SNr neuron maximum firing rates from the input-output curve (WT: N = 6, n = 64; PNKD: N = 7, n = 101; Mann-Whitney U test, \*\*\*\**P* < 0.0001). Boxplots show median interquartile range (IQR) and 1.5 IQR (b,c,g,k,o,p). Data shown as mean ± SEM (n).

Next, we examined action potential waveforms, which reflect ion channel composition and kinetics. To evoke action potentials under similar conditions, we held SNr neurons at -70 mV to silence their spontaneous activity and standardize the availability of voltage-gated sodium channels across WT and PNKD neurons. We then evoked spiking with a 500 ms, 500 pA current step and analyzed the resulting action potentials in two ways: using the first evoked action potential (Fig. 6d–g) or the average of all action potentials evoked across the sweep (Fig. 6h–k). While action potential shapes were qualitatively similar between WT and PNKD mice, we found the spike height, which tends to reflect voltage-gated sodium conductance, was lower in PNKD SNr neurons (Fig. 6e,g). The properties of voltage-gated sodium channels can also be captured by plotting the action potential phase plot, the first derivative of voltage (dV/dt) versus the absolute voltage (Fig. 6f). In SNr neurons from PNKD mice, both the peak dV/dt and maximal voltage excursion are reduced, suggesting a reduction in sodium channel density. These changes in spike properties were even more pronounced when analyzed across an entire sweep (Fig. 6h–k). Other properties, including threshold and afterhyperpolarization amplitude, did not differ between genotypes, although the first evoked action potential was slightly narrower in PNKD neurons (Table S2). Overall, these findings are consistent with a decrease in sodium channel availability in PNKD versus WT SNr neurons.

To assess the excitability of SNr neurons, we injected a series of increasing amplitude current steps and recorded the resulting firing rate of each neuron. In WT neurons, current injections elicited progressively higher firing rates (Fig. 6l, left). By contrast, PNKD neurons “stuttered” at lower amplitude current injections and entered depolarization block at current injections that elicited sustained firing in WT neurons (Fig. 6l, right). Maximum firing rates were shifted toward lower values in SNr neurons from PNKD compared to WT mice (Fig. 6m). Notably, nearly a quarter of PNKD SNr neurons were unable to fire consistent action potentials at any current injection. Even when comparing sweeps in which SNr neurons could fire across a step, neurons from PNKD mice fired at lower rates than those from WT mice (Fig. 6n). PNKD neurons also had a higher rheobase (Fig. 6o) and attained a lower maximum firing rate (Fig. 6p). Taken together, PNKD SNr neurons show several alterations in their intrinsic properties, most notably lower maximum firing rates.

### Striatal MSN activity is reduced during dystonic attacks

We next sought to define what triggers depolarization block *in vivo*. In PNKD mice, many SNr neurons transiently increased firing after ethanol administration, just before the loss of firing (Fig. 1i and Fig. 3a). Increased firing can lead to inactivation of voltage-gated sodium channels (Na_v_), causing depolarization block. What circuit mechanisms might drive this initial increase in SNr firing? SNr neurons receive highly convergent inhibitory inputs from GABAergic D1-expressing striatal medium spiny neurons (D1-MSNs)^61,62^. Thus, a reduction in D1-MSN activity is well positioned to disinhibit (increase) SNr firing. To test whether ethanol decreases D1-MSN firing in PNKD mice, we recorded single-unit activity from MSNs in the sensorimotor, dorsolateral striatum (DLS) of WT and PNKD mice before and after ethanol injection. In a subset of recordings, we identified D1-MSNs with an opto-tagging protocol. In these recordings, we injected an AAV encoding Cre-dependent ChR2-eYFP and implanted an optrode in the DLS of WT and PNKD;D1-Cre mice (Fig. 7a; Fig. S3a–b). Optically identified cells showed consistent short-latency spiking in response to blue light pulses (Fig. 7b). We first examined the activity of all MSNs before and after ethanol injection. In contrast to recordings in the SNr, MSN firing rates were comparable in WT and PNKD mice during the baseline period (Fig. 7c–d; Fig. S3c– d). In WT mice, the firing of MSNs fell slightly overall after ethanol injection (Fig. S3c–d). By contrast, the firing rate of MSNs markedly decreased during the post-ethanol period in PNKD mice (Fig. 7c,d). We then examined the D1-MSN subpopulation. In WT mice, D1-MSN firing did not systematically change after ethanol injection (Fig. S3e,f). However, in PNKD mice, we observed a consistent decrease in D1-MSN firing after ethanol administration (Fig. 7e,f). These results are consistent with the idea that ethanol-induced loss of striatal inhibition of SNr neurons trigger dystonic attacks.

**Fig. 7:**
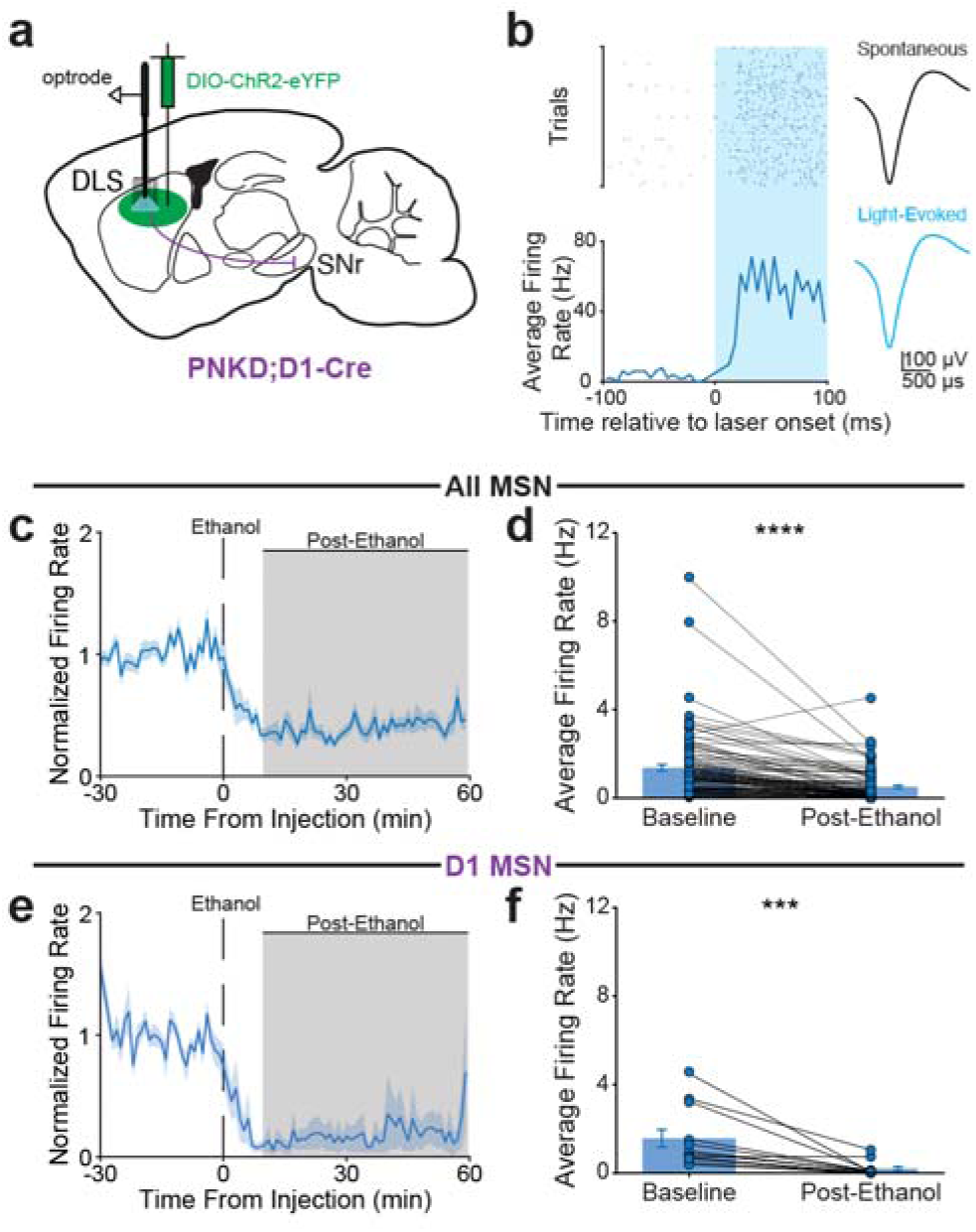
Striatal MSN activity is reduced during dystonic attacks. **a,** Schematic showing optrode recording configuration and injection of AAV encoding DIO-ChR2-eYFP in the dorsal lateral striatum (DLS) of PNKD;D1-Cre or PNKD;A2a-Cre mice. **b,** Representative optogenetically-labeled medium spiny neuron (MSN), showing responses to blue light stimulation (left) and average spontaneous and light-evoked waveforms (right). **c,** Average normalized firing rate of all MSNs before and after ethanol injection in PNKD mice. **d,** Comparison of MSN average firing rates during baseline (0–30 minutes prior to injection) and post-ethanol (10–60 minutes after injection) periods (N = 9, n = 107, Wilcoxon signed-rank test, \*\*\*\**P* < 0.0001). **e,** Average normalized firing rate of D1-MSNs before and after ethanol injection in PNKD mice. **f,** Comparison of D1-MSN average firing rates during baseline and post-ethanol periods (N = 4, n = 12, Wilcoxon signed-rank test, \*\*\**P* = 0.0005).

### Pulsatile activation of inhibitory striatal inputs to the SNr attenuates dystonia

Our data demonstrated dramatically reduced SNr firing during dystonic attacks, and suggested synaptic disinhibition followed by depolarization block as a candidate mechanism. To test the causal role of these phenomena in dystonia, we attempted to hyperpolarize SNr neurons to relieve depolarization block *in vivo*. We injected AAV encoding ChR2-eYFP in the dorsal striatum (DS) of PNKD mice and implanted optical fibers bilaterally over the SNr, allowing us to optically activate the inhibitory striatal projections that terminate there (Fig. 8a–c). In control animals, we injected an AAV encoding the inert fluorophore mCherry in the DS. To validate this approach, we optically stimulated striatal terminals in acute SNr slices from PNKD mice (Fig. 8d). In whole-cell current-clamp mode, we drove SNr neurons into depolarization block with a sustained 10-second current injection and, on alternating sweeps, delivered pulsatile blue light (4 Hz, 8 ms pulses, 0.5 mW) to activate striatal terminals (Fig. 8e–j). We alternated light-off and light-on sweeps (3 each, counterbalanced) for a within cell comparison of the effect of light stimulation (averaged from 1-10 sec). As expected, in slices from control mice, light had no effect and SNr neurons remained silent (Fig. 8e–g). In slices from ChR2-expressing mice, light pulses were hyperpolarizing and were followed by rebound spiking suggestive of recovery from depolarization block (Fig. 8h,i). On average, brief optical stimulation of DS fibers markedly increased the net firing rate of SNr neurons (Fig. 8j). These experiments suggest that pulsatile optical stimulation of inhibitory inputs to SNr neurons promote recovery of spiking in PNKD mice.

**Fig. 8:**
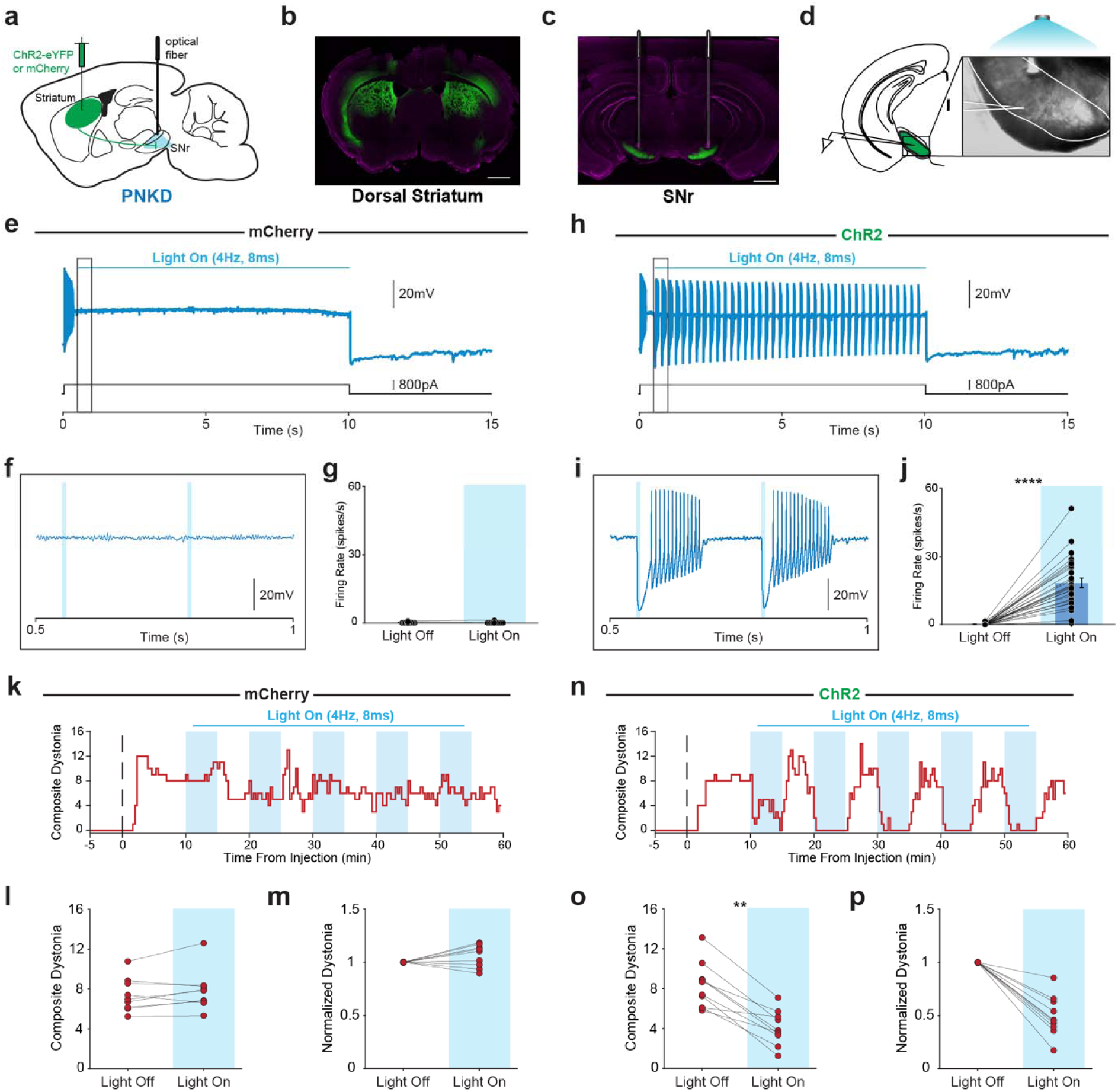
Pulsatile activation of inhibitory striatal inputs to the SNr attenuates dystonia. **a,** Schematic showing bilateral injection of an AAV encoding ChR2-eYFP or mCherry in the dorsal striatum and implantation of optical fiber above the SNr in PNKD animals. **b,** Representative postmortem confirmation of viral expression in the dorsal striatum. White scale bar represents 1 mm. **c,** Representative postmortem confirmation of striatal projections to the SNr and optical fiber placement above the SNr. White scale bar represents 1 mm. **d,** Schematic (left) and DIC image (right) of whole cell, current clamp electrophysiological recording configuration. **e–j,** Recordings from SNr neurons in mice expressing mCherry (e–g) or ChR2-eYFP (h–j) in the dorsal striatum. To test whether striatal input can relieve depolarization block, cells were first driven into depolarization block with a 10 second step current injection. Beginning 550 ms into the step, blue light (4 Hz, 8 ms pulses, 0.5 mW) was delivered to activate striatal terminals. Light-off (laser was turned off) and light-on sweeps were alternated across six repetitions, and firing rate was averaged over 1–10 seconds of each sweep. **e,** Representative light-on sweep in a mouse that expressed mCherry. **f,** Expanded view of the sweep in (e) to show detail. **g,** Comparison of firing rate during light-off and light-on sweeps in mice that expressed mCherry (N = 2, n = 11), Wilcoxon signed-rank test, *P* = 0.8750). **h,** Representative light-on sweep in a mouse that expressed ChR2. **i,** Expanded view of the sweep in (h) to show detail. **j,** Comparison of firing rate during light-off and light-on sweeps in mice that expressed ChR2 (N = 4, n = 27, Wilcoxon signed-rank test, \*\*\*\**P* < 0.0001). **k–p**, *In vivo* dystonia severity in mice expressing mCherry (k–m) or ChR2-eYFP (n–p) in the dorsal striatum. 10 minutes after ethanol injection, each mouse received alternating light-off and light-on epochs (five each), with the starting condition counterbalanced across mice. Light-on epochs consisted of bilateral blue light stimulation of the SNr at 4 Hz, 8 ms pulses, and 3–6 mW. **k,** Representative time course of ethanol induced dystonia severity in a mCherry expressing PNKD mouse with alternating light epochs (light-on periods shaded blue). **l,** Average composite dystonia during light-off and light-on conditions (N = 9, Wilcoxon signed-rank test, *P* = 0.1641) in mCherry-expressing mice. **m,** Average dystonia normalized to light off severity for mCherry-expressing mice. **n**, Representative time course of ethanol induced dystonia severity in a ChR2 expressing PNKD mouse with alternating light epochs (light-on periods shaded blue). **o,** Average composite dystonia during light-off and light-on periods (N = 10, Wilcoxon signed-rank test, \*\**P* = 0.0095) in ChR2-expressing mice. **p,** Average dystonia normalized to light-off severity for ChR2-expressing mice. Data shown as mean ± SEM (g,j)

We tested whether such stimulation could be effective *in vivo* to attenuate dystonic attacks. In the same PNKD mice described above, we induced dystonia with ethanol and then delivered blue light pulses in alternating light-off and light-on epochs (Fig. S4), using the same patterned stimulation used in slice experiments. Both ChR2-expressing and mCherry-expressing mice were split into two groups, counterbalancing which epoch came first to control for changes in dystonia over the session (Fig. 8k–p; Fig. S4). In mCherry-expressing mice, SNr light stimulation had no effect on dystonia severity (Fig. 8k–m). By contrast, in ChR2-expressing mice, dystonia severity was significantly reduced during light-on epochs (Fig. 8n–p; Supplementary Video 2). On average, dystonia was reduced by ∼50% (Fig. 8p) and this effect was reversible, as it closely tracked the light-on and light-off epochs across the session (Fig. 8n; Fig. S4). Together, these results demonstrate that restoring SNr firing during dystonic attacks by allowing neurons to recover from depolarization block is sufficient to reduce dystonia.

## Discussion

Our study tested the hypothesis that reduced SNr activity underlies dystonia. Using the PNKD mouse model, we found that SNr firing was profoundly reduced during dystonic attacks. Several lines of evidence indicated that SNr neurons enter depolarization block during dystonia, including broader and lower amplitude single-unit spike waveforms prior to cessation of firing, and increased intracellular calcium during dystonic attacks, as measured by GCaMP fiber photometry. Moreover, during dystonic attacks, ChR2-expressing SNr neurons were unable to spike in response to optogenetic stimulation. We found that ethanol triggers decreased D1-MSN firing, which may lead to transient disinhibition of SNr neurons, facilitating depolarization block in vulnerable PNKD SNr neurons with limited maximum firing rates. Finally, relieving depolarization block with pulsatile optogenetic activation of striatal inhibitory inputs restored SNr firing and attenuated dystonia. These results suggest that PNKD dystonia is driven by depolarization block in SNr neurons.

The finding that SNr activity is reduced during dystonia is consistent with decades of human studies implicating the basal ganglia in the disorder. Dystonia occurs in a diverse range of syndromes with many etiologies^1,63,64^. However, this etiological heterogeneity may converge on a shared set of motor pathways, raising the possibility that reduced activity in basal ganglia output nuclei is a final common pathway of dystonia rather than a mechanism unique to PNKD. Consistent with this idea, many (though not all) forms of dystonia respond to basal ganglia deep brain stimulation (DBS). As in most dystonias, individuals with PNKD mutations show no overt neurodegeneration, suggesting the disorder arises from aberrant neural activity or connectivity. Human intraoperative recordings during DBS implantation or pallidotomy have similarly found lower firing rates in basal ganglia output nuclei compared to healthy non-human primates or individuals with Parkinson’s disease^22–26^. These recordings in the GPi have also identified increased bursting of single units and low-frequency (2–10 Hz) oscillatory activity in the local field potential that may reflect a pathological neural rhythm in dystonia^24,65^. Unfortunately, there are significant limitations in the interpretation of human physiology in people with dystonia, including the use of anesthesia and challenges correlating neurophysiology with dystonia on shorter timescales. In addition, human studies currently cannot establish cell-type specificity nor causal relationships between neural activity and dystonia. Our results address this gap by demonstrating both a temporal correlation and a causal relationship between SNr firing and dystonic attacks in a mouse model.

While we examined basal ganglia activity in the PNKD model, human dystonia has been linked to multiple brain regions and circuits, including the basal ganglia, cerebellum and cortex. A newer hypothesis posits that dystonia may result from dysfunction in one or multiple nodes that triggers aberrant signals throughout a network^48,66–71^. For example, a primary process in the cerebellum or basal ganglia may both result in abnormal activity in classical sensorimotor pathways, resulting in dystonia. Altered physiology may propagate between the cerebellum and basal ganglia, via direct or indirect connections, including midbrain dopamine neurons^72–78^.

Most rodent models of dystonia involve a trade-off between recapitulating the dystonic phenotype and mirroring the molecular origins of human dystonia. Transgenic rodent models of common forms of inherited dystonia do not often develop frank dystonia. For example, mice carrying the *TOR1A* mutation that causes human DYT1 dystonia develop only minor motor abnormalities^31,32^. Other animal models do develop dystonia, but rely on pharmacology, spontaneous mutations or genetic manipulations not seen in humans, and thus have limitations in terms of explaining the pathophysiology of dystonia. Nonetheless, these phenotypic models provide valuable evidence that dysfunction in the basal ganglia^79–81^, cerebellum^36–38,82–85^, or forebrain^86^ *can* produce dystonia, at least in rodents. PNKD mice are one of the few mouse models that is both genotypic and phenotypic (also see Chiken et al^79^). While the rarity and paroxysmal nature of PNKD may limit the generalizability of our findings^43,44,47^, the model’s phenotypic penetrance may relate to its paroxysmal nature: brief attacks leave less time for homeostatic compensation. Discrete dystonic episodes also allow comparison of SNr activity within animals, revealing markedly lower rates during dystonia versus healthy movement.

Multiple lines of evidence suggest that SNr neurons enter depolarization block, leading to profoundly suppressed firing during dystonic attacks in PNKD mice. While *in vivo* extracellular electrophysiology has limitations in establishing depolarization block as a mechanism, our *ex vivo* intracellular recordings corroborate altered intrinsic properties and a vulnerability to depolarization block in SNr neurons from PNKD mice. What prevents depolarization block under normal conditions and what triggers depolarization block when PNKD mice are exposed to ethanol? PNKD is expressed broadly in neurons^43^, and mutations in the *PNKD* gene may cause a primary disruption in cellular and synaptic function, which in turn leads to homeostatic changes in excitability and/or synaptic connectivity, helping people and mice with PNKD mutations to function normally under almost all circumstances. Stimuli such as intense stress, caffeine, and ethanol may push neuronal activity outside this compensated range, leading to a breakdown in physiological function.

Depolarization block results from accumulated Na_v_ inactivation, often due to an increase in firing. Consistent with this idea, we found that in many SNr neurons, firing transiently increased after ethanol administration, coinciding with a decrease in the activity of D1-MSNs, the predominant inhibitory input to the SNr^61^. In PNKD mice, the decrease in D1-MSN activity may be sufficient to drive transient increases in SNr firing and tip neurons into depolarization block, and the mouse into dystonia. Interestingly, we found highly consistent decreases in D1-MSN firing in PNKD mice treated with ethanol; in a prior study we found more variable changes in D1-MSN firing when mice were treated with caffeine, which in turn triggers hyperkinetic attacks with only transient, if any, dystonia^42,41^. Differences in both behavioral phenotype and D1-MSN activity between caffeine and ethanol may be a clue as to the role of D1-MSN activity in dystonia versus other hyperkinetic movements. We cannot rule out that increased excitatory drive from the subthalamic nucleus (STN) contributes to triggering SNr depolarization block in PNKD mice. However, the overall decrease in MSN activity (including D2-MSNs) in response to ethanol would be expected to reduce STN activity via the indirect pathway^87,88^. Isolating the pharmacological mechanism by which ethanol triggers circuit dysfunction in PNKD is difficult, as it acts on a wide range of targets throughout the brain^89^. At doses similar to those used here, ethanol increases dopamine release and inhibits NMDA-type glutamate receptors in the striatum^90^. In a previous study, PNKD mice treated with caffeine showed elevated levels of dopamine metabolites, as compared to WT mice^41^. Here, acute ethanol had little effect on MSN or SNr activity in WT mice. This result suggests that cellular or circuit changes in PNKD mice, possibly dysfunctional ethanol metabolism, dopamine signaling or altered synaptic connectivity, may trigger transient increases in SNr activity not seen in the healthy brain.

The susceptibility of SNr neurons to depolarization block may stem from changes in intrinsic properties. Healthy GABAergic SNr neurons fire at high tonic rates driven by tightly coordinated ion channels^49,50,91^. Between spikes, a persistent Na_v_ current and tonically active TRPC3 current slowly depolarize the membrane toward threshold^51,92^. At threshold, a high-density transient Na_v_ current drives the rising phase of the action potential and triggers voltage-dependent K^+^ channels to rapidly repolarize the membrane^93–95^. This intrinsic firing mechanism appears impaired in PNKD SNr neurons. Even without ethanol, these neurons fired at lower rates than WT in both *in vivo* and *ex vivo* recordings. In PNKD (as compared to WT) SNr neurons, action potentials had lower spike height, implying reduced Na_v_ availability. Interestingly, the average membrane potentials of PNKD SNr neurons in whole-cell configuration were slightly more hyperpolarized than WT potentials, suggesting multiple channels beyond the spike-generating Na_v_ channels may be affected. Our *in vivo* physiology recordings suggest that PV-SNr neurons are especially vulnerable to depolarization block. These cells are particularly high-firing, and their vulnerability may be due to disruption of their unique ion channel composition^49,50,96^.

What might drive reduced Na channel availability in PNKD SNr neurons? While the normal function of the PNKD protein (formerly MR-1) remains unknown, the protein is expressed in neurons and may modulate synaptic release^97,98^. However, PNKD is likely mediated by gain-of-function mutations^40,45^. The molecular consequences of these mutations are unclear, leaving open whether reduced Na channel availability reflects a direct effect of PNKD dysfunction or a homeostatic response. This apparent vulnerability does not generalize across cell types, as striatal MSNs from PNKD mice have intrinsic properties similar to WT neurons^42^. Indeed, striatal neurons in PNKD mice show alterations in synaptic function and synaptic plasticity^42^. Regardless of mechanism, these baseline changes alone are not sufficient to trigger dystonia, as PNKD mice remain asymptomatic until challenged with a trigger such as ethanol.

Ultimately, the goal of this work is to inform the development of therapies that help people living with dystonia. Our optogenetic rescue established that restoring SNr firing is sufficient to attenuate dystonia. Though optogenetics is not currently feasible in the human basal ganglia, our work suggests a clear physiological target. Pharmacological or gene therapy approaches targeting ion channel function in basal ganglia output nuclei may be an alternative strategy.

## Methods

### Animals

All experimental procedures were approved by the University California, San Francisco Institutional Animal Care and Use Committee. Hemizygous PNKD mice (Tg(RP24-112K19 Pnkd*/IRESDsRed)), hereafter termed “PNKD mice”), were maintained on a C57Bl/6 background. For all experiments, hemizygous PNKD mice were bred to wild-type (WT) C57Bl/6 (Jackson Laboratory #000664) or hemizygous Parvalbumin-Cre (PV-Cre; Jackson Laboratory #017320), Adora2a-Cre (A2a-Cre, Jackson Laboratory #040186), or Drd1-Cre (D1-Cre; Jackson Laboratory #040188). PNKD-negative littermates were used as WT controls. We used 3–7 month old male and female mice. Animals were housed under a 12h light/dark cycle with access to food and water ad libitum.

### Behavior

Details of blinded dystonia assessment can be found at dx.doi.org/10.17504/protocols.io.j8nlk7b9wg5r/v2. Before all *in vivo* experiments, mice were habituated to handling and injections by receiving daily intraperitoneal (i.p.) saline for a minimum of 1 week. They were also habituated to the open field chamber for a minimum of 1 hour daily for 2 days prior to experiments. During experiments, mice were placed in the open field chamber on top of a clear acrylic elevated stage. Mice were monitored by two cameras (The Imaging Source): one mounted directly below the clear acrylic stage (bottom-up view) and one mounted to the side of the chamber (side view). After a baseline of 30 minutes, animals were injected IP with either saline or 1.5 g/kg ethanol in saline. Dystonia was scored offline using both bottom-up and side camera views. The experimenter was blinded to animal name, genotype, and injection type. Dystonia was scored across four body segments: (1) trunk/neck, (2) forelimb, (3) hindlimb, and (4) tail. The severity of dystonia was quantified using a time-based scale ranging from 0–4 (0 = 0%, 1 = <50%, 2 = 51%-80%, 3 = 81%-95%, 4 = 95%-100%). A composite dystonia score was calculated by adding dystonia severity across each body segment for an overall score ranging 0–16. The dystonia score was calculated across time bins ranging 1–5 minutes in length.

### Surgical procedures

A detailed surgery protocol can be found at DOI: https://dx.doi.org/10.17504/protocols.io.b9kxr4xn. Briefly, mice were anesthetized with a combination of ketamine/xylazine (40/10 mg/kg IP) and inhaled isoflurane (1%). When unresponsive to a toe pinch, mice were placed on a heating pad and head-fixed in a stereotactic frame (Kopf Instruments). Lubricant with ophthalmic ointment was applied to the eyes. At the incision site, fur was removed with Nair and alternating 70% ethanol and betadine was applied to clean the scalp before an incision was made. All burr holes and craniectomies were made with a mounted drill. Virus was delivered into the brain using a 33-gauge needle (WPI) and Micro4 pump (WPI).

For *in vivo* single-unit electrophysiology with optical identification in the SNr, PV-Cre mice were stereotaxically injected with AAV5-DIO-ChR2-eYFP (1:3 dilution, 500nL, Addgene v40448) into the SNr (-3.2 AP, -1.6 ML, -4.5 DV).C For optical identification experiments in the DLS, D1-Cre or A2a-Cre mice were injected with AAV5-DIO-ChR2-eYFP (1:2 dilution, 1 µl, UPenn Vector Core) into the DLS (+0.8 AP, -2.25 ML, -2.5 DV). To construct optrodes, optical fiber-ferrule assemblies were cemented onto either a microelectrode array (custom 16-channel, 28-channel, or 32-channel; Innovative Neurophysiology), or a high density 64 channel silicon probe (Cambridge Neurotech H6, 2 shanks, 9 mm length). Optical fiber-ferrule assemblies consisted of a 200 µm optical fiber (Thorlabs) threaded through a ceramic ferrule (Thorlabs) and cemented with epoxy. The tip of the optic fiber was positioned ∼0.5 mm above the tip of the electrode array or in the middle of the silicon probe contacts. To implant electrophysiological recording devices, the animal underwent a separate surgery 4 weeks after viral injection. For implantation of the microelectrode array, the device was slowly lowered through a craniectomy into the left SNr or DLS and secured with a thin layer of dental cement (Metabond) and dental acrylic (Henry Schein). A ground wire was inserted into the right cerebellum and covered with dental cement and acrylic. Experiments were performed between one week and four weeks after implantation. For silicon probe implantation, a drivable 2-shank 64-channel high density silicon probe (model H6, Cambridge Neurotech) was mounted on a movable Microdrive and the recording electrodes were stained with DiO lipophilic dye (ThermoFisher). The probe was then implanted above the SNr (-3.5 DV) and secured with dental cement and acrylic. A ground wire was also inserted into the right cerebellum and secured. At least three days prior to the probe implant, a metal head bar and empty Microdrive were attached to skull with dental cement and acrylic to habituate the animal to added weight. Experiments were performed between one day and two weeks after implantation.

For fiber photometry experiments, PV-Cre mice were injected with AAV1-DIO-GCaMP6s (1:9 dilution, 500nL, UPenn P2824) into the SNr (-4.5 DV) and implanted with a fiber-optic ferrule (0.4 mm, Doric Lenses) above the SNr (-4.2 DV). Fibers were secured with dental cement and dental acrylic. Experiments were performed at least 4 weeks after surgery. For *ex vivo* slice electrophysiology experiments, 2–3-month-old PV-Cre mice were injected with AAV5-DIO-mCherry (undiluted, 500nL, UNC AV4634C) into the SNr. Slice experiments were performed 4– 5 weeks after surgery. For optogenetic stimulation of striatal inputs to SNr, PNKD mice were injected with either AAV5-hSyn-ChR2-eYFP (1:8 dilution, Addgene v191297) or AAV-hSyn-mCherry (undiluted, Vectorbuilder 211112AAVD16). 500 nL of virus was injected into 4 sites on each hemisphere to cover the dorsal striatum (AP: 0.8, ML: ±2.2, DV:-2.5; AP: 0.8, ML: ±1.2, DV:-2.5; AP: 0.0, ML: ±2.0, DV:-2.5; AP: 0.5, ML: ±2.5, DV:-2.0). In the same surgery, optical fibers (400 µm, Doric Lenses) were implanted bilaterally above the SNr (-4.2 DV) and secured with dental cement and acrylic. Experiments were performed at least 4 weeks after surgery.

### In vivo electrophysiology

A detailed protocol for *in vivo* electrophysiology can be found here: https://dx.doi.org/10.17504/protocols.io.36wgq641ylk5/v1. Prior to experimental sessions, animals were habituated to daily i.p. injections for a week and at least two 1-hour sessions in the open field chamber (clear acrylic, 25 cm diameter). *In vivo* single unit electrophysiological recordings were made using two different recording devices: custom fixed microwire (35 mm tungsten) arrays (16-channel, 28 channel, 32-channel; Innovative Neurophysiology) or a drivable high-density 64-channel silicon probe (Cambridge Neurotech H6, 2 shanks, 9 mm length). For recordings using microelectrode arrays, mice were habituated to i.p. injection, tethering, and the open arena for a minimum of 1 hour on 2 separate sessions. Recordings began 1-2 weeks after implantation. For recordings using a silicon probe, animals were habituated to tethering and the open arena for a minimum of 1 hour. Recordings began a minimum of 1 day after implantation and lasted for no longer than 2 weeks. During SNr recordings, animals were tethered to a lightweight, ultra-thin SPI interface cable (INTAN) attached to an electrical commutator (Doric). For mice implanted with a microelectrode array, the device was attached to the SPI cable via a RHD 32-channel recording head stage (INTAN).

During striatal recordings, animals were tethered to a lightweight, multiplexed headstage cable (Dragonfly/Triangle Biosystems). SNr electrical signals were acquired at 30 kHz using an INTAN system (RHD2000 USB Interface Board, INTAN). Simultaneously, video was taken by TTL-triggered bottom-up and side view cameras at 25 fps using IC Capture software. The same TTL pulse used to trigger the cameras was also sent to the INTAN recording system to synchronize video and electrophysiology signal. Ethanol injection was synchronized with a manually triggered TTL pulse. Striatal electrical signals were recorded on a MAP system, using RASPUTIN 2.4 HLK3 acquisition software (Plexon).

Recording sessions consisted of a 30-minute baseline period followed by an injection of ethanol. If a mouse developed any involuntary movements during the baseline period, the experiment was terminated. Recordings lasted 1.5–6 hours total. For sessions that involved optogenetic cell identification, an experimenter began observing the mice after 1-hour post-injection. Once the animal showed no abnormal movements for 10 consecutive minutes, the animal was tethered to a patch cord (Doric Lenses) connected to a blue laser (Shanghai Laser and Optics Century) via an optical commutator (Doric Lenses). The laser was controlled by TTL pulses delivered from an Arduino. The identification protocol consisted of 500 ms blue light pulses delivered at 2 Hz for 2 minutes in four different light powers, including 0.5, 1, 2, and 4 mW. For experiments determining if optical activation during an attack was sufficient to increase spiking in the SNr, this same optical protocol was applied 10–20 minutes after ethanol injection and then again upon behavioral recovery. For visualization and analysis across baseline and post-ethanol periods, the firing rates of single units were averaged in 1-minute bins.

Single units from recordings in the SNr were identified using automated spike sorting algorithms (KiloSort, https://github.com/cortex-lab/Kilosort). After sorting, single units were manually curated in Phy based on waveform (to remove clusters with axonal characteristics), waveform amplitude drift, and ISI distribution. Single units from striatal recordings were identified by manual sorting in Offline Sorter 3.3.5 (Plexon). Waveform features used for separating units were typically a combination of valley amplitude, the first principal components, and/or nonlinear energy. Clusters were classified as single units if they fulfilled the following criteria: (1) the cluster was statistically different from multiunit activity and from other single-unit clusters on the same wire (*P* < 0.05, MANOVA), and (2) no interspike interval <1 ms observed. Single units were classified as putative MSNs based on waveform and interspike interval distribution, using previously published criteria to exclude fast-spiking interneurons^99^.

To optically identify PV-SNr and D1-MSN single units, spike times were aligned to laser onsets offline. A threshold was set at the 99% confidence interval of the pre-light baseline (50 ms before light on). PV-expressing SNr units were classified as optically identified if they met all three of the following criteria: (1) firing rate exceeded threshold within 5 ms of laser onset; (2) firing rate exceeded threshold for at least 20 ms; and (3) light-evoked spike waveforms were not distinguishable from spontaneous waveforms. MSN units were classified as optically identified by the same criteria except firing rate had to exceed threshold within 15 ms of laser onset.

To extract and compare waveforms in our *in vivo* physiology data, we adapted code originally contributed by C. Shoonover and A. Fink (found here: https://github.com/cortex-lab/spikes/blob/master/analysis/getWaveForms.m). Baseline waveforms were obtained by randomly sampling 20 waveforms 10–5 minutes before ethanol injection. For recordings in PNKD animals, we first identified the time at which firing rate fell below the 99% CI of baseline firing or the onset of a period of ≥ 5 seconds without an action potential. We then sampled 20 waveforms in the 5-second window directly preceding that time. For recordings in WT animals, we sampled 20 waveforms from a randomly selected 5-second window that fell within 2–6 minutes after ethanol injection. This time range was chosen, as rate changes occurred within 2–6 minutes after ethanol injection in PNKD recordings. The amplitude and width of each waveform was calculated and then averaged to compare across time periods. Amplitude was defined as the change in voltage from the onset of the spike to the peak. Width was defined as the time between spike onset and peak.

### Fiber photometry

A detailed fiber photometry protocol can be found here: https://dx.doi.org/10.17504/protocols.io.b9udr6s6. Prior to experimental sessions, animals were habituated to daily i.p. injections for a week and at least two 1-hour sessions in the open field chamber (clear acrylic, 25 cm diameter). On testing days, mice were placed in the open field arena where GCaMP signals and videos were recorded for a 20-minute photobleaching period, a 20-minute baseline, and for at least 1 hour after saline or ethanol injection. Fiber photometry signals were acquired through surgically implanted 400 µm optical fibers coupled to an LED driver system (Doric). Following signal modulation, 405 nm (isosbestic) and 465 nm (calcium-dependent) signals were demodulated via a lock-in amplifier (RZ5P, TDT), visualized, and recorded (Synapse, TDT). Videos were taken by TTL-triggered bottom-up and side view cameras at 25 fps using IC Capture software. The same TTL pulses used to trigger the cameras were also sent to the photometry recoding system to synchronize video and GCAMP signal. Ethanol injection was marked in the photometry acquisition file with a manually triggered TTL pulse.

Fiber photometry signals were demodulated from the raw TDT stream and analyzed in MATLAB. To correct for calcium-independent artifacts including motion, the 405 nm signal was linearly fit to the 465 nm signal using a first-degree polynomial. The fitted coefficients were then applied to the full isosbestic trace to generate a scaled reference signal F_0_. Fluorescent signals (dF/F) were calculated for each sample as 1 + (F_465_ – F_0_) / F_0_. The resulting dF/F signals were visualized in two ways: (1) by converting dF/F to z-scores using the mean and standard deviation of the entire recording session or (2) by normalizing dF/F to the average baseline dF/F. Statistical comparison was conducted using dF/F.

### Ex vivo slice electrophysiology

A detailed protocol for *ex vivo* electrophysiology can be found here: dx.doi.org/10.17504/protocols.io.b9uir6ue. All animals were handled and injected i.p. with saline for 5 days a week for a minimum of two weeks prior to slice recordings to minimize the possibility of stress-induced dyskinetic attacks at the time of sacrifice^41^. To prepare acute brain slices, mice were deeply anesthetized with a ketamine/xylazine injection and transcardially perfused with carbogenated, ice-cold glycerol-based artificial cerebrospinal fluid (ACSF) containing (in mM): 250 glycerol, 2.5 KCl, 1.2 NaH_2_PO_4_, 10 HEPES, 21 NaHCO_3_, 5 D-Glucose, 2 MgCl_2_, 2 CaCl_2_. After decapitation, the brain was removed and mounted to a submerged chuck in a slicing chamber containing ice-cold, carbogenated glycerol-based ACSF where coronal slices (275μm) containing the SNr were collected with a vibratome (Leica). Slices were immediately transferred to a chamber containing warm (33°–34°C), carbogenated ACSF containing the following (in mM): 125 NaCl, 26 NaHCO_3_, 2.5 KCl, 1 MgCl_2_, 2 CaCl_2_, 1.25 NaH_2_PO_4_, 12.5 glucose for 30-60 min, then stored in carbogenated ACSF at room temperature. All slice recordings occurred in a chamber superfused (∼2 mL/min) with carbogenated, warm (29°–32°C) ACSF. Neurons were patched using borosilicate glass electrodes (2-5 MΩ) filled with potassium-based internal solution containing the following (in mM): 130 KMeSO_3_, 10 NaCl, 2 MgCl_2_, 0.16 CaCl_2_, 0.5 EGTA, 10 HEPES, 2 MgATP, 0.3 NaGTP, (pH 7.3). Neurons were recorded in cell-attached and whole-cell configuration using a MultiClamp 700B amplifier (Molecular Devices) and digitized with an ITC-18 A/D board (HEKA). Data were acquired using Igor Pro 6.0 software (Wavemetrics) and custom acquisition routines (mafPC, courtesy of M.A. Xu-Friedman). Recordings were acquired at 20 kHz and filtered at 2 kHz.

For experiments to measure the intrinsic properties of SNr neurons, PV-SNr neurons were identified by their mCherry-positive somata. Spontaneous activity of the cells was first recorded in cell-attached mode and then in whole-cell current-clamp mode. Immediately after whole-cell break in, action potential shapes were collected in current clamp by holding the cell at -70 mV and then delivering a 500 pA, 500 ms current injection. Spike shapes were compared in two ways: using the first evoked action potential or the average of all action potentials evoked across the sweep. The spike threshold was defined as the membrane voltage at the first time point at which dV/dt exceeded 25 V/s. The spike amplitude was measured as the difference between the peak voltage of the waveform and the threshold voltage. Spike width was measured by taking the difference between the time points at which the voltage trace crossed threshold (filtered to exclude crossings separated by fewer than 6 samples to suppress noise-driven crossings). Lastly, the spike afterhyperpolarization (AHP) amplitude was defined as the minimum membrane voltage that occurred within 5 ms of peak voltage. Whole-cell spontaneous firing rate and average membrane potential were recorded in current clamp. To calculate the input resistance of each cell, the recording was switched to voltage clamp, the cell was held at -70 mV and a series of -5 pA current steps were delivered; the resulting change in voltage was averaged across sessions. Current-clamp recordings were then made to obtain the input-output properties of SNr neurons. A series of 450 ms square-wave current steps ranging from 10 pA to 4000 pA were delivered until the cell was driven into depolarization block. The maximum firing rate was obtained by calculating spikes/s during the current injection on the last sweep across which the cell could sustain firing. If a cell did not fire consistent action potentials at any current injection, it was assigned a maximum firing rate of 0 Hz.

For experiments to validate optical activation of striatal synaptic inputs to the SNr, and determine whether period activation of these inhibitory inputs could restore firing after depolarization block, SNr neurons were recorded in whole-cell configuration. To determine the current injection that would reliably induce depolarization block in each cell, we used the same protocol of square-wave current steps described above. Using that current step, SNr neurons were driven into depolarization block with a sustained 10-second current injection, and on alternating sweeps, pulsatile blue light was delivered (4 Hz, 8 ms pulses, 0.5 mW). Light-off and light-on sweeps were alternated (3 each, counterbalanced) for a within-cell comparison of the effect of light stimulation (averaged spikes/s from 1-10 sec).

### Optogenetics

Optogenetics was used in several components of this study. For a description of its use in optical labeling of single-unit electrophysiology, see the *In Vivo* Electrophysiology section above. For a description of its use in *ex vivo* electrophysiology, see the *Ex Vivo* Slice Electrophysiology section above. This section is devoted to describing its use for *in vivo* circuit manipulation during dystonia (Figure 8). A detailed protocol for optogenetic manipulations can be found here: https://dx.doi.org/10.17504/protocols.io.b9wfr7bn. To optogenetically stimulate striatal inputs to the SNr, PNKD mice were injected with either AAV5-hSyn-ChR2-eYFP or AAV5-hSyn-mCherry (control) into the dorsal striatum and implanted with 400 µm fibers above the SNr (see Surgical Procedures). Prior to experimental sessions, animals were habituated to daily IP injections for a week and at least two 1-hour sessions in the open field chamber (clear acrylic, 25 cm diameter). To account for mouse-to-mouse variability in the ability to activate inhibitory inputs to the SNr (ChR2 expression, fiber placement), before experimental sessions, we identified an optimally therapeutic stimulation power in mice expressing ChR2. Animals were tethered to a lightweight patch cable (Doric Lenses) connected via optical commutator (Doric Lenses) to a blue laser (Shanghai Laser) and placed in the open field chamber. Dystonia was induced by i.p. ethanol administration (1.5mg/kg). The laser and cameras were controlled via TTL pulses from an Arduino Uno and all TTLs were recorded in Intan software. At peak dystonia, we blue light (4Hz, 8ms) was delivered at a range of powers 0.5–6 mW. The power at which dystonia was most attenuated was used for later experimental sessions.

Each session began with a 5-minute baseline period, followed by ethanol administration and a 10-minute period without stimulation. Subsequently, 5 light-on (blue light, 4 Hz, 8 ms pulses, 1– 6 mW) and 5 light off epochs (5 min each) were alternated. To control for changes in dystonia severity over time, mice were randomly assigned before the session to one of two counterbalanced groups: Group A began with a light-on epoch 10 minutes after ethanol injection, whereas Group B began with a light-off epoch (first light-on epoch at 15 minutes). Sessions were recorded with side and bottom-up video and aligned offline to the times of IP ethanol injection and optical stimulation epochs. Each video session was then split into 5-minute clips spanning baseline through the final light-on or light-off epoch. Clips were given random filenames and scored in random order, leaving the rater with no information about the injected virus (ChR2 vs mCherry) or the clip’s position within the session. To quantify the effect of stimulation, each mouse’s dystonia score was averaged across its five light-on epochs and its five light-off epochs.

### Histology and microscopy

For *in vivo* electrophysiology experiments, the location of each recorded single unit was confirmed to be within the SNr (units falling outside the SNr were excluded). To verify the location of channels on microelectrode arrays, prior to terminal transcardial perfusion, animals were deeply anesthetized and electrolytic lesions were made by applying 100 uA for 5 seconds per each microwire. To verify the location of channels on high-density probes, DiI signal (applied at the time of surgical implantation) was histologically registered to the Allen Mouse Brain Atlas (https://github.com/cortex-lab/allenCCF^100^). Following all *in vivo* experiments, mice were deeply anesthetized with a ketamine/xylazine injection and transcardially perfused with 4% PFA in PBS. The brain was then dissected out, postfixed in PFA overnight, and transferred to 30% sucrose at 4°C for cryoprotection. Brains were cut on a freezing microtome (Leica) into 30–40 µm sections and stored in PBS. Following *ex vivo* electrophysiology experiments, 275 μm slices were fixed overnight in 4% PFA. The next day, the tissue was transferred to 30% sucrose and re-sectioned into 50 μm slices using a freezing microtome (Leica). All sections were mounted in Vectashield Mounting Medium with DAPI on glass slides for subsequent imaging. Images (4X and 20X) were obtained using an Olympus Fluoview FV3000 confocal microscope or a Nikon Ti Inverted microscope.

### Experimental design and statistical analysis

The experimental design and statistical analysis for all key experiments are summarized in Table S1. Where stated, sample sizes for both animals (N) and cells (n) were pre-determined based on a power calculation using pilot data or existing literature. We aimed to achieve >90% power to detect significance with a two-sided α of 0.05, using statistical tests reported in Table S1. *Ex vivo* electrophysiology traces were processed in Igor Pro 6.3 (Wavemetrics). Statistical tests were performed in MATLAB.

## Data and Code availability

Data and original code has been deposited at Zenodo (DOI: 10.5281/zenodo.21998092).

## Acknowledgements

This work was funded by NSF GRFP 2445150 to OKB and R01 NS 131276 to ABN. We thank Kevin Bender, Louis Ptáček, Ying-Hui Fu, and members of the Nelson Laboratory for their valuable feedback on the manuscript.

## References

1. Albanese A, Bhatia K, Bressman SB, et al. Phenomenology and classification of dystonia: A consensus update. 10.1002/mds.25475. Movement Disorders. 2013/06/15 2013;28(7):863-873. 10.1002/mds.25475

2. Shakkottai VG, Batla A, Bhatia K, et al. Current Opinions and Areas of Consensus on the Role of the Cerebellum in Dystonia. Cerebellum. Apr 2017;16(2):577–594. doi:10.1007/s12311-016-0825-6

3. Tewari A, Fremont R, Khodakhah K. It’s not just the basal ganglia: Cerebellum as a target for dystonia therapeutics. Mov Disord. Nov 2017;32(11):1537–1545. doi:10.1002/mds.27123

4. Albin RL, Young AB, Penney JB. The functional anatomy of basal ganglia disorders. Trends in Neurosciences. 1989/01/01/ 1989;12(10):366-375. 10.1016/0166-2236(89)90074-X

5. DeLong MR. Primate models of movement disorders of basal ganglia origin. Trends in Neurosciences. 1990/07/01/ 1990;13(7):281-285. 10.1016/0166-2236(90)90110-V

6. Mink JW. The Basal Ganglia and Involuntary Movements: Impaired Inhibition of Competing Motor Patterns. Archives of Neurology. 2003;60(10):1365–1368. doi:10.1001/archneur.60.10.1365

7. Vitek JL. Pathophysiology of dystonia: a neuronal model. Mov Disord. 2002;17 Suppl 3:S49–62. doi:10.1002/mds.10142

8. Bhatia KP, Marsden CD. The behavioural and motor consequences of focal lesions of the basal ganglia in man. Brain. Aug 1994;117 ( Pt 4):859–76. doi:10.1093/brain/117.4.859

9. Marsden CD, Obeso JA, Zarranz JJ, Lang AE. The anatomical basis of symptomatic hemidystonia. Brain. Jun 1985;108 ( Pt 2):463–83. doi:10.1093/brain/108.2.463

10. Ostrem JL, Starr PA. Treatment of dystonia with deep brain stimulation. Neurotherapeutics. Apr 2008;5(2):320–30. doi:10.1016/j.nurt.2008.01.002

11. FitzGerald JJ, Rosendal F, de Pennington N, et al. Long-term outcome of deep brain stimulation in generalised dystonia: a series of 60 cases. J Neurol Neurosurg Psychiatry. Dec 2014;85(12):1371–6. doi:10.1136/jnnp-2013-306833

12. Kupsch A, Benecke R, Müller J, et al. Pallidal deep-brain stimulation in primary generalized or segmental dystonia. N Engl J Med. Nov 9 2006;355(19):1978–90. doi:10.1056/NEJMoa063618

13. Wichmann T, DeLong MR. Deep Brain Stimulation for Movement Disorders of Basal Ganglia Origin: Restoring Function or Functionality? Neurotherapeutics. Apr 2016;13(2):264–83. doi:10.1007/s13311-016-0426-6

14. Mink JW. THE BASAL GANGLIA: FOCUSED SELECTION AND INHIBITION OF COMPETING MOTOR PROGRAMS. Progress in Neurobiology. 1996/11/01/ 1996;50(4):381-425. 10.1016/S0301-0082(96)00042-1

15. Hikosaka O. Basal ganglia--possible role in motor coordination and learning. Curr Opin Neurobiol. Dec 1991;1(4):638–43. doi:10.1016/s0959-4388(05)80042-x

16. Hikosaka O. Neural systems for control of voluntary action--a hypothesis. Adv Biophys. 1998;35:81–102.

17. Mink JW, Thach WT. Basal ganglia intrinsic circuits and their role in behavior. Curr Opin Neurobiol. Dec 1993;3(6):950–7. doi:10.1016/0959-4388(93)90167-w

18. Nelson AB, Kreitzer AC. Reassessing models of basal ganglia function and dysfunction. Annu Rev Neurosci. 2014;37:117–35. doi:10.1146/annurev-neuro-071013-013916

19. Wichmann T, DeLong MR. Models of basal ganglia function and pathophysiology of movement disorders. Neurosurg Clin N Am. Apr 1998;9(2):223–36.

20. Berardelli A, Rothwell JC, Hallett M, Thompson PD, Manfredi M, Marsden CD. The pathophysiology of primary dystonia. Brain. Jul 1998;121 ( Pt 7):1195–212. doi:10.1093/brain/121.7.1195

21. Kaymak A, Colucci F, Ahmadipour M, et al. Spiking Patterns in the Globus Pallidus Highlight Convergent Neural Dynamics across Diverse Genetic Dystonia Syndromes. Ann Neurol. May 2025;97(5):826–844. doi:10.1002/ana.27185

22. Sanghera MK, Grossman RG, Kalhorn CG, Hamilton WJ, Ondo WG, Jankovic J. Basal ganglia neuronal discharge in primary and secondary dystonia in patients undergoing pallidotomy. Neurosurgery. Jun 2003;52(6):1358–70; discussion 1370-3. doi:10.1227/01.neu.0000064805.91249.f5

23. Vitek JL, Chockkan V, Zhang JY, et al. Neuronal activity in the basal ganglia in patients with generalized dystonia and hemiballismus. Ann Neurol. Jul 1999;46(1):22–35. doi:10.1002/1531-8249(199907)46:1<22::aid-ana6>3.0.co;2-z

24. Starr PA, Rau GM, Davis V, et al. Spontaneous pallidal neuronal activity in human dystonia: comparison with Parkinson’s disease and normal macaque. J Neurophysiol. Jun 2005;93(6):3165–76. doi:10.1152/jn.00971.2004

25. Tang JK, Moro E, Mahant N, et al. Neuronal firing rates and patterns in the globus pallidus internus of patients with cervical dystonia differ from those with Parkinson’s disease. J Neurophysiol. Aug 2007;98(2):720–9. doi:10.1152/jn.01107.2006

26. Sedov A, Dzhalagoniya I, Semenova U, et al. Unraveling the neural signatures: Distinct pallidal patterns in dystonia subtypes. Parkinsonism Relat Disord. Jan 2025;130:107207. doi:10.1016/j.parkreldis.2024.107207

27. Tanabe LM, Martin C, Dauer WT. Genetic background modulates the phenotype of a mouse model of DYT1 dystonia. PLoS One. 2012;7(2):e32245. doi:10.1371/journal.pone.0032245

28. Sharma N, Baxter MG, Petravicz J, et al. Impaired motor learning in mice expressing torsinA with the DYT1 dystonia mutation. J Neurosci. Jun 1 2005;25(22):5351–5. doi:10.1523/jneurosci.0855-05.2005

29. Grundmann K, Reischmann B, Vanhoutte G, et al. Overexpression of human wildtype torsinA and human DeltaGAG torsinA in a transgenic mouse model causes phenotypic abnormalities. Neurobiol Dis. Aug 2007;27(2):190–206. doi:10.1016/j.nbd.2007.04.015

30. Shashidharan P, Sandu D, Potla U, et al. Transgenic mouse model of early-onset DYT1 dystonia. Hum Mol Genet. Jan 1 2005;14(1):125–33. doi:10.1093/hmg/ddi012

31. Dang MT, Yokoi F, McNaught KS, et al. Generation and characterization of Dyt1 DeltaGAG knock-in mouse as a model for early-onset dystonia. Exp Neurol. Dec 2005;196(2):452–63. doi:10.1016/j.expneurol.2005.08.025

32. Goodchild RE, Kim CE, Dauer WT. Loss of the dystonia-associated protein torsinA selectively disrupts the neuronal nuclear envelope. Neuron. Dec 22 2005;48(6):923–32. doi:10.1016/j.neuron.2005.11.010

33. Hodge AT, Rasheed MA, Lin T, Burgess CR, Leventhal DK. Cortically Dependent Motor Training Does Not Induce Abnormal Movements in DYT1-Knock In Mice. Brain and Behavior. 2026;16(1):e71176. 10.1002/brb3.71176

34. Richter A, Löscher W. Pathophysiology of idiopathic dystonia: findings from genetic animal models. Progress in neurobiology. 1998;54(6):633–677.

35. Lorden J, McKeon TW, Baker H, Cox N, Walkley S. Characterization of the rat mutant dystonic (dt): a new animal model of dystonia musculorum deformans. The Journal of neuroscience. 1984;4(8):1925–1932.

36. Brown AM, van der Heijden ME, Jinnah HA, Sillitoe RV. Cerebellar Dysfunction as a Source of Dystonic Phenotypes in Mice. Cerebellum. Aug 2023;22(4):719–729. doi:10.1007/s12311-022-01441-0

37. White JJ, Sillitoe RV. Genetic silencing of olivocerebellar synapses causes dystonia-like behaviour in mice. Nat Commun. Apr 4 2017;8:14912. doi:10.1038/ncomms14912

38. Calderon DP, Fremont R, Kraenzlin F, Khodakhah K. The neural substrates of rapid-onset Dystonia-Parkinsonism. Nat Neurosci. Mar 2011;14(3):357–65. doi:10.1038/nn.2753

39. Jinnah HA, Hess EJ, Ledoux MS, Sharma N, Baxter MG, Delong MR. Rodent models for dystonia research: characteristics, evaluation, and utility. Mov Disord. Mar 2005;20(3):283–92. doi:10.1002/mds.20364

40. Shen Y, Lee HY, Rawson J, et al. Mutations in PNKD causing paroxysmal dyskinesia alters protein cleavage and stability. Hum Mol Genet. Jun 15 2011;20(12):2322–32. doi:10.1093/hmg/ddr125

41. Lee H-y, Nakayama J, Xu Y, et al. Dopamine dysregulation in a mouse model of paroxysmal nonkinesigenic dyskinesia. The Journal of Clinical Investigation. 02/01/ 2012;122(2):507-518. doi:10.1172/JCI58470

42. Nelson AB, Girasole AE, Lee HY, Ptáček LJ, Kreitzer AC. Striatal Indirect Pathway Dysfunction Underlies Motor Deficits in a Mouse Model of Paroxysmal Dyskinesia. J Neurosci. Mar 30 2022;42(13):2835–2848. doi:10.1523/jneurosci.1614-20.2022

43. Lee HY, Xu Y, Huang Y, et al. The gene for paroxysmal non-kinesigenic dyskinesia encodes an enzyme in a stress response pathway. Hum Mol Genet. Dec 15 2004;13(24):3161–70. doi:10.1093/hmg/ddh330

44. Pons R, Cuenca-León E, Miravet E, et al. Paroxysmal non-kinesigenic dyskinesia due to a PNKD recurrent mutation: report of two Southern European families. Eur J Paediatr Neurol. Jan 2012;16(1):86–9. doi:10.1016/j.ejpn.2011.09.008

45. Shen Y, Ge W-P, Li Y, et al. Protein mutated in paroxysmal dyskinesia interacts with the active zone protein RIM and suppresses synaptic vesicle exocytosis. Proceedings of the National Academy of Sciences. 2015/03/10 2015;112(10):2935-2941. doi:10.1073/pnas.1501364112

46. Bruno MK, Lee HY, Auburger GW, et al. Genotype-phenotype correlation of paroxysmal nonkinesigenic dyskinesia. Neurology. May 22 2007;68(21):1782–9. doi:10.1212/01.wnl.0000262029.91552.e0

47. Erro R, Sheerin U-M, Bhatia KP. Paroxysmal dyskinesias revisited: A review of 500 genetically proven cases and a new classification. Movement Disorders. 2014/08/01 2014;29(9):1108–1116. 10.1002/mds.25933

48. Corp DT, Greenwood CJ, Morrison-Ham J, et al. Clinical and Structural Findings in Patients With Lesion-Induced Dystonia: Descriptive and Quantitative Analysis of Published Cases. Neurology. Nov 1 2022;99(18):e1957–e1967. doi:10.1212/wnl.0000000000201042

49. Delgado-Zabalza L, Mallet NP, Glangetas C, et al. Targeting parvalbumin-expressing neurons in the substantia nigra pars reticulata restores motor function in parkinsonian mice. Cell Rep. Oct 31 2023;42(10):113287. doi:10.1016/j.celrep.2023.113287

50. McElvain LE, Chen Y, Moore JD, et al. Specific populations of basal ganglia output neurons target distinct brain stem areas while collateralizing throughout the diencephalon. Neuron. May 19 2021;109(10):1721–1738.e4. doi:10.1016/j.neuron.2021.03.017

51. Atherton JF, Bevan MD. Ionic mechanisms underlying autonomous action potential generation in the somata and dendrites of GABAergic substantia nigra pars reticulata neurons in vitro. J Neurosci. Sep 7 2005;25(36):8272–81. doi:10.1523/jneurosci.1475-05.2005

52. Jin X, Costa RM. Start/stop signals emerge in nigrostriatal circuits during sequence learning. Nature. Jul 22 2010;466(7305):457–62. doi:10.1038/nature09263

53. González-Hernández T, Rodríguez M. Compartmental organization and chemical profile of dopaminergic and GABAergic neurons in the substantia nigra of the rat. Journal of Comparative Neurology. 2000/05/22 2000;421(1):107-135. 10.1002/(SICI)1096-9861(20000522)421:1<107::AID-CNE7>3.0.CO;2-F

54. Lee CR, Tepper JM. Morphological and physiological properties of parvalbumin- and calretinin-containing γ-aminobutyric acidergic neurons in the substantia nigra. Journal of Comparative Neurology. 2007/02/10 2007;500(5):958-972. 10.1002/cne.21220

55. Vilin YY, Ruben PC. Slow inactivation in voltage-gated sodium channels: molecular substrates and contributions to channelopathies. Cell Biochem Biophys. 2001;35(2):171–90. doi:10.1385/cbb:35:2:171

56. Rudy B. Slow inactivation of the sodium conductance in squid giant axons. Pronase resistance. J Physiol. Oct 1978;283:1–21. doi:10.1113/jphysiol.1978.sp012485

57. Fleidervish IA, Friedman A, Gutnick MJ. Slow inactivation of Na+ current and slow cumulative spike adaptation in mouse and guinea-pig neocortical neurones in slices. J Physiol. May 15 1996;493 ( Pt 1)(Pt 1):83–97. doi:10.1113/jphysiol.1996.sp021366

58. Milner ES, Do MTH. A Population Representation of Absolute Light Intensity in the Mammalian Retina. Cell. Nov 2 2017;171(4):865–876.e16. doi:10.1016/j.cell.2017.09.005

59. Hodgkin AL, Huxley AF. A quantitative description of membrane current and its application to conduction and excitation in nerve. J Physiol. Aug 1952;117(4):500–44. doi:10.1113/jphysiol.1952.sp004764

60. Thaler C, Gray AC, Lipscombe D. Cumulative inactivation of N-type CaV2.2 calcium channels modified by alternative splicing. Proc Natl Acad Sci U S A. Apr 13 2004;101(15):5675–9. doi:10.1073/pnas.0303402101

61. Deng YP, Lei WL, Reiner A. Differential perikaryal localization in rats of D1 and D2 dopamine receptors on striatal projection neuron types identified by retrograde labeling. J Chem Neuroanat. Dec 2006;32(2-4):101–16. doi:10.1016/j.jchemneu.2006.07.001

62. Foster NN, Barry J, Korobkova L, et al. The mouse cortico–basal ganglia– thalamic network. Nature. 2021/10/01 2021;598(7879):188-194. doi:10.1038/s41586-021-03993-3

63. Prudente CN, Hess EJ, Jinnah HA. Dystonia as a network disorder: what is the role of the cerebellum? Neuroscience. Feb 28 2014;260:23–35. doi:10.1016/j.neuroscience.2013.11.062

64. Neychev VK, Gross RE, Lehéricy S, Hess EJ, Jinnah HA. The functional neuroanatomy of dystonia. Neurobiol Dis. May 2011;42(2):185–201. doi:10.1016/j.nbd.2011.01.026

65. Silberstein P, Kühn AA, Kupsch A, et al. Patterning of globus pallidus local field potentials differs between Parkinson’s disease and dystonia. Brain. Dec 2003;126(Pt 12):2597–608. doi:10.1093/brain/awg267

66. Argyelan M, Carbon M, Niethammer M, et al. Cerebellothalamocortical connectivity regulates penetrance in dystonia. J Neurosci. Aug 5 2009;29(31):9740–7. doi:10.1523/jneurosci.2300-09.2009

67. Niethammer M, Carbon M, Argyelan M, Eidelberg D. Hereditary dystonia as a neurodevelopmental circuit disorder: Evidence from neuroimaging. Neurobiol Dis. May 2011;42(2):202–9. doi:10.1016/j.nbd.2010.10.010

68. Shakkottai VG. Physiologic changes associated with cerebellar dystonia. Cerebellum. Oct 2014;13(5):637–44. doi:10.1007/s12311-014-0572-5

69. Corp DT, Joutsa J, Darby RR, et al. Network localization of cervical dystonia based on causal brain lesions. Brain. Jun 1 2019;142(6):1660–1674. doi:10.1093/brain/awz112

70. Jinnah HA, Hess EJ. Evolving concepts in the pathogenesis of dystonia. Parkinsonism & Related Disorders. 2018/01/01/ 2018;46:S62-S65. 10.1016/j.parkreldis.2017.08.001

71. Quartarone A, Hallett M. Emerging concepts in the physiological basis of dystonia. Mov Disord. Jun 15 2013;28(7):958–67. doi:10.1002/mds.25532

72. Person AL, Gale SD, Farries MA, Perkel DJ. Organization of the songbird basal ganglia, including area X. J Comp Neurol. Jun 10 2008;508(5):840–66. doi:10.1002/cne.21699

73. Ichinohe N, Mori F, Shoumura K. A di-synaptic projection from the lateral cerebellar nucleus to the laterodorsal part of the striatum via the central lateral nucleus of the thalamus in the rat. Brain Res. Oct 13 2000;880(1-2):191–7. doi:10.1016/s0006-8993(00)02744-x

74. Hoshi E, Tremblay L, Féger J, Carras PL, Strick PL. The cerebellum communicates with the basal ganglia. Nat Neurosci. Nov 2005;8(11):1491–3. doi:10.1038/nn1544

75. Bostan AC, Dum RP, Strick PL. The basal ganglia communicate with the cerebellum. Proc Natl Acad Sci U S A. May 4 2010;107(18):8452–6. doi:10.1073/pnas.1000496107

76. Watabe-Uchida M, Zhu L, Ogawa SK, Vamanrao A, Uchida N. Whole-brain mapping of direct inputs to midbrain dopamine neurons. Neuron. Jun 7 2012;74(5):858–73. doi:10.1016/j.neuron.2012.03.017

77. Chen CH, Fremont R, Arteaga-Bracho EE, Khodakhah K. Short latency cerebellar modulation of the basal ganglia. Nat Neurosci. Dec 2014;17(12):1767–75. doi:10.1038/nn.3868

78. Washburn S, Oñate M, Yoshida J, et al. The cerebellum directly modulates the substantia nigra dopaminergic activity. Nat Neurosci. Mar 2024;27(3):497–513. doi:10.1038/s41593-023-01560-9

79. Chiken S, Shashidharan P, Nambu A. Cortically evoked long-lasting inhibition of pallidal neurons in a transgenic mouse model of dystonia. J Neurosci. Dec 17 2008;28(51):13967–77. doi:10.1523/jneurosci.3834-08.2008

80. Bennay M, Gernert M, Richter A. Spontaneous remission of paroxysmal dystonia coincides with normalization of entopeduncular activity in dt (SZ) mutants. The Journal of neuroscience: the official journal of the Society for Neuroscience. 2001;21(13):RC153-RC153.

81. Gernert M, Bennay M, Fedrowitz M, Rehders JH, Richter A. Altered discharge pattern of basal ganglia output neurons in an animal model of idiopathic dystonia. Journal of Neuroscience. 2002;22(16):7244–7253.

82. Pizoli CE, Jinnah HA, Billingsley ML, Hess EJ. Abnormal cerebellar signaling induces dystonia in mice. J Neurosci. Sep 1 2002;22(17):7825–33. doi:10.1523/jneurosci.22-17-07825.2002

83. Fremont R, Calderon DP, Maleki S, Khodakhah K. Abnormal high-frequency burst firing of cerebellar neurons in rapid-onset dystonia-parkinsonism. J Neurosci. Aug 27 2014;34(35):11723–32. doi:10.1523/jneurosci.1409-14.2014

84. Fremont R, Tewari A, Khodakhah K. Aberrant Purkinje cell activity is the cause of dystonia in a shRNA-based mouse model of Rapid Onset Dystonia-Parkinsonism. Neurobiol Dis. Oct 2015;82:200–212. doi:10.1016/j.nbd.2015.06.004

85. Washburn S, Fremont R, Moreno-Escobar MC, Angueyra C, Khodakhah K. Acute cerebellar knockdown of Sgce reproduces salient features of myoclonus-dystonia (DYT11) in mice. Elife. Dec 23 2019;8doi:10.7554/eLife.52101

86. Kernodle K, Bakerian AM, Cropsey A, Dauer WT, Leventhal DK. A dystonia mouse model with motor and sequencing deficits paralleling human disease. Behavioural Brain Research. 2022/05/24/ 2022;426:113844. 10.1016/j.bbr.2022.113844

87. Alexander GE, DeLong MR, Strick PL. Parallel organization of functionally segregated circuits linking basal ganglia and cortex. Annu Rev Neurosci. 1986;9:357–81. doi:10.1146/annurev.ne.09.030186.002041

88. Alexander GE, Crutcher MD. Functional architecture of basal ganglia circuits: neural substrates of parallel processing. Trends Neurosci. Jul 1990;13(7):266–71. doi:10.1016/0166-2236(90)90107-l

89. Lovinger DM, Roberto M. Synaptic effects induced by alcohol. Curr Top Behav Neurosci. 2013;13:31–86. doi:10.1007/7854_2011_143

90. Lovinger DM, Alvarez VA. Alcohol and basal ganglia circuitry: Animal models. Neuropharmacology. Aug 1 2017;122:46–55. doi:10.1016/j.neuropharm.2017.03.023

91. Zhou FM, Lee CR. Intrinsic and integrative properties of substantia nigra pars reticulata neurons. Neuroscience. Dec 15 2011;198:69–94. doi:10.1016/j.neuroscience.2011.07.061

92. Zhou FW, Matta SG, Zhou FM. Constitutively active TRPC3 channels regulate basal ganglia output neurons. J Neurosci. Jan 9 2008;28(2):473–82. doi:10.1523/jneurosci.3978-07.2008

93. Hille B. Sinauer Associates; Sunderland, MA: 2001. Ion Channels of Excitable Membranes[Google Scholar].

94. Bean BP. The action potential in mammalian central neurons. Nature Reviews Neuroscience. 2007/06/01 2007;8(6):451-465. doi:10.1038/nrn2148

95. Ding S, Matta SG, Zhou FM. Kv3-like potassium channels are required for sustained high-frequency firing in basal ganglia output neurons. J Neurophysiol. Feb 2011;105(2):554–70. doi:10.1152/jn.00707.2010

96. Oh SW, Harris JA, Ng L, et al. A mesoscale connectome of the mouse brain. Nature. 2014/04/01 2014;508(7495):207–214. doi:10.1038/nature13186

97. Shen Y, Ge WP, Li Y, et al. Protein mutated in paroxysmal dyskinesia interacts with the active zone protein RIM and suppresses synaptic vesicle exocytosis. Proc Natl Acad Sci U S A. Mar 10 2015;112(10):2935–41. doi:10.1073/pnas.1501364112

98. Ghezzi D, Viscomi C, Ferlini A, et al. Paroxysmal non-kinesigenic dyskinesia is caused by mutations of the MR-1 mitochondrial targeting sequence. Hum Mol Genet. Mar 15 2009;18(6):1058–64. doi:10.1093/hmg/ddn441

99. Gage GJ, Stoetzner CR, Wiltschko AB, Berke JD. Selective activation of striatal fast-spiking interneurons during choice execution. Neuron. Aug 12 2010;67(3):466–79. doi:10.1016/j.neuron.2010.06.034

100. Shamash P, Carandini M, Harris K, Steinmetz N. A tool for analyzing electrode tracks from slice histology. bioRxiv. 2018:447995. doi:10.1101/447995

