## Supplementary Figures and Tables for "Depolarization block in substantia nigra pars reticulata neurons drives dystonia in a mouse model"

**Extended Data**

**Table S1.** Experimental Design and analysis*^a^*

| **Figure** | **Parameter** | **Data Type** | **Statistical Test** | **Planned N(mice), n(cells)** | **Actual N(mice), n(cells)** | **Mean ± SEM** | **Statistical Result** |
| --- | --- | --- | --- | --- | --- | --- | --- |
| 1c,d | Average Dystonia | WT | MWU | N.A. | N = 12 | 6.69 ± 0.67 | *P* < 0.0001 |
|  |  | PNKD |  | N.A. | N = 12 | 0 ± 0 |  |
| 1h | Average Firing Rate (Hz) WT | Baseline | WSR | n = 50 | N = 7 n = 75 | 32.90 ± 2.84 | *P* = 0.0395 |
|  |  | Post-Ethanol |  |  |  | 30.13 ± 2.66 |  |
| 1k | Average Firing Rate (Hz) PNKD | Baseline | WSR | n = 50 | N = 6 n = 46 | 21.32 ± 3.50 | *P* < 0.0001 |
|  |  | Post-Ethanol |  |  |  | 6.78 ± 2.80 |  |
| 1h,k | Average Baseline Firing Rate (Hz) | WT | MWU | n = 50 | N = 7 n = 75 | 32.90 ± 2.84 | *P* = 0.0029 |
|  |  | PNKD |  | n = 50 | N = 6 n = 46 | 21.32 ± 3.50 |  |
| S2a | Time to Rate ∆ (Per-Cell) vs Time to Dystonia | PNKD | Pearson | N.A. | N = 6 n = 44 | N.A. | *r = 0.471 R^2^ = 0.222 P = 0.0013* |
| S2a | Time to Rate ∆ (Per-Recording) vs Time to Dystonia | PNKD | Pearson | N.A. | N = 6 n = 11 | N.A. | *r = 0.760 R^2^ = 0.578 P = 0.0066* |
| S2b | Average Firing Rate (Hz) PNKD | Baseline | repeated measures ANOVA | n = 20 | N = 5 n = 34 | 25.6 ± 4.46 | *F*(2,66) = 12.377  *P* < 0.0001 |
|  |  | Post-Ethanol |  |  |  | 6.59 ± 3.58 |  |
|  |  | Recovery |  |  |  | 20.64 ± 2.30 |  |
| S2b | Average Firing Rate (Hz) PNKD | Baseline | Post Hoc WSR | n = 50 | N = 5 n = 34 | 25.6 ± 4.46 | *P* < 0.0001 |
|  |  | Post-Ethanol |  |  |  | 6.59 ± 3.58 |  |
|  |  | Baseline |  |  |  | 25.6 ± 4.46 | *P* = 0.7518 |
|  |  | Recovery |  |  |  | 20.64 ± 2.30 |  |
|  |  | Post-Ethanol |  |  |  | 6.59 ± 3.58 | *P* < 0.0001 |
|  |  | Recovery |  |  |  | 20.64 ± 2.30 |  |
| 2d | Average Firing Rate (Hz) WT;PV+ | Baseline | WSR | n = 20 | N = 3 n = 21 | 40.40 ± 6.00 | *P* = 0.6639 |
|  |  | Post-Ethanol |  |  |  | 38.18 ± 5.50 |  |
| 2f | Average Firing Rate (Hz) PNKD;PV+ | Baseline | WSR | n = 20 | N = 4 n = 16 | 11.35 ± 2.16 | *P* < 0.0004 |
|  |  | Post-Ethanol |  |  |  | 0.24 ± 0.10 |  |
| 2d,f | Average Baseline Firing Rate PV+ | WT | MWU | n = 20 | N = 3 n = 21 | 40.40 ± 6.00 | *P* < 0.0001 |
|  |  | PNKD |  |  | N = 4 n = 16 | 11.35 ± 2.16 |  |
| 3c | Spike Amplitude (µV) PNKD | Baseline | WSR | N.A. | N = 6 n = 40 | 394.14 ± 33.51 | *P* < 0.0001 |
|  |  | Pre-Rate Decrease |  |  |  | 296.92 ± 19.32 |  |
| 3d | Spike Width (ms) PNKD | Baseline | WSR | N.A. | N = 6 n = 40 | 0.25 ± 0.02 | *P* < 0.0001 |
|  |  | Pre-Rate Decrease |  |  |  | 0.35 ± 0.03 |  |
| 3g | Spike Amplitude (µV) WT | Baseline | WSR | N.A. | N = 7 n = 75 | 276.65 ± 13.46 | *P* = 0.3195 |
|  |  | Matched Time Window |  |  |  | 279.44 ± 12.80 |  |
| 3h | Spike Width (ms) WT | Baseline | WSR | N.A. | N = 7 n = 75 | 0.26 ± 0.02 | *P* = 0.7453 |
|  |  | Matched Time Window |  |  |  | 0.26 ± 0.01 |  |
| 4d | Average dF/F WT | Baseline | WSR | N = 8 | N = 9 | 1.03 ± 0.01 | *P* = 0.0039 |
|  |  | Post-Ethanol |  |  |  | 0.99 ± 0.01 |  |
| 4f | Average dF/F PNKD | Baseline | WSR | N = 8 | N = 7 | 0.91 ± 0.02 | *P* = 0.0156 |
|  |  | Post-Ethanol |  |  |  | 1.13 ± 0.04 |  |
| 5e | Average Firing Rate (Hz) Recovered PNKD | Pre-Light | WSR | n = 10 | N = 5 n = 16 | 28.66 ± 4.59 | *P* = 0.0004 |
|  |  | Light On |  |  |  | 113.06 ± 16.64 |  |
| 5f | Average Firing Rate (Hz) During Dystonia PNKD | Pre-Light | WSR |  |  | 0.55 ± 0.44 | *P* = 0.625 |
|  |  | Light On |  |  |  | 0.52 ± 0.42 |  |
| 6b | Cell Attached Firing Rate (Hz) | WT | MWU | N.A. | N = 6 n = 58 | 15.67 ± 1.49 | *P* = 0.0002 |
|  |  | PNKD |  |  | N = 7 n = 63 | 11.09 ± 1.49 |  |
| 6c | Whole Cell Firing Rate (Hz) | WT | MWU | N.A. | N = 6 n = 62 | 16.78 ± 2.42 | *P* = 0.0188 |
|  |  | PNKD |  |  | N = 7 n = 32 | 10.34 ± 1.81 |  |
| 6f | First Action Potential  Peak dvdt (V/s) | WT | MWU | n = 70 | N = 6 n = 69 | 586.47 ± 12.82 | *P* = 0.0033 |
|  |  | PNKD |  | n = 70 | N = 7 n = 96 | 531.87 ± 13.01 |  |
| 6g | First Action Potential Spike Height (mV) | WT | MWU | n = 70 | N = 6 n = 69 | 82.51 ± 0.01 | *P* < 0.0001 |
|  |  | PNKD |  | n = 70 | N = 7 n = 96 | 74.78 ± 0.01 |  |
| 6j | Average Action Potential Peak dvdt (V/s) | WT | MWU | n = 70 | N = 6 n = 69 | 349.02 ± 12.98 | *P* = 0.0006 |
|  |  | PNKD |  | n = 70 | N = 7 n = 96 | 284.11 ± 12.46 |  |
| 6k | Average Action Potential Spike Height (mV) | WT | MWU | n = 70 | N = 6 n = 69 | 60.43 ± 1.11 | *P* < 0.0001 |
|  |  | PNKD |  | n = 70 | N = 7 n = 96 | 50.75 ± 1.06 |  |
| 6m | Survivor Trace | WT | KS | n = 70 | N = 6 n = 64 | N.A. | *D =* 0.367  *P* < 0.0001 |
|  |  | PNKD |  | n = 70 | N = 7 n = 101 | N.A. |  |
| 6n | Input-Output Curve | WT | LMM | n = 70 | N = 6 n = 63 | N.A. | current: F(1,2039) = 3789.00  *P* < 0.0001  genotype: F(1,2039) = 21.46  *P* < 0.0001  cxg: F(1,2039) = 21.98  *P* < 0.0001 |
|  |  | PNKD |  | n = 70 | N = 7 n = 77 | N.A. |  |
| 6o | Rheobase (pA) | WT | MWU | n = 70 | N = 6 n = 62 | 10.48 ± 4.33 | *P* < 0.0001 |
|  |  | PNKD |  | n = 70 | N = 7 n = 77 | 86.75 ± 14.92 |  |
| 6p | Maximum Firing Rate (Hz) | WT | MWU | n = 70 | N = 6 n = 64 | 284.42 ± 10.38 | *P* < 0.0001 |
|  |  | PNKD |  | n = 70 | N = 7 n = 101 | 180.75 ± 10.54 |  |
| 7d | Average Firing Rate (Hz) PNKD;All MSNs | Baseline | WSR | N.A. | N = 9 n = 107 | 1.40 ± 0.14 | *P* < 0.0001 |
|  |  | Post-Ethanol |  |  |  | 0.48 ± 0.07 |  |
| 7f | Average Firing Rate (Hz) PNKD;D1+ | Baseline | WSR | N.A. | N = 4 n = 12 | 1.56 ± 0.40 | *P* = 0.0005 |
|  |  | Post-Ethanol |  |  |  | 0.19 ± 0.10 |  |
| S3d | Average Firing Rate (Hz) WT;All MSNs | Baseline | WSR | N.A. | N = 10 n = 139 | 1.71 ± 0.20 | *P* = 0.0015 |
|  |  | Post-Ethanol |  |  |  | 1.46 ± 0.22 |  |
| S3f | Average Firing Rate (Hz) WT;D1+ | Baseline | WSR | N.A. | N = 2 n = 17 | 1.28 ± 0.24 | *P* = 0.0615 |
|  |  | Post-Ethanol |  |  |  | 1.03 ± 0.18 |  |
| 8g | Average Firing Rate (Hz) PNKD, mCherry | Light Off | WSR | N = 3 n = 15 | N = 2 n = 11 | 0.13 ± 0.09 | *P* = 0.8750 |
|  |  | Light On |  |  |  | 0.11 ± 0.11 |  |
| 8j | Average Firing Rate (Hz) PNKD, ChR2 | Light Off | WSR | N = 3 n = 15 | N = 4 n = 27 | 0.17 ± 0.09 | *P < 0.0001* |
|  |  | Light On |  |  |  | 18.38 ± 2.13 |  |
| 8m | Average Dystonia PNKD, mCherry | Light Off | WSR | N = 8 | N = 9 | 7.42 ± 0.57 | *P* = 0.1641 |
|  |  | Light On |  |  |  | 7.86 ± 0.68 |  |
| 8o | Average Dystonia PNKD, ChR2 | Light Off | WSR | N = 8 | N = 10 | 8.28 ± 0.73 | *P* = 0.0095 |
|  |  | Light On |  |  |  | 4.06 ± 0.54 |  |

*^a^*MWU: Mann-Whitney *U* test; WSR: Wilcoxon signed-rank test; KS: two-sample Kolmogorov-Smirnov test; LMM: linear mixed-effects model.

**Supplementary Video 1. Motor behavior in PNKD mouse before and after ethanol injection.** This video shows two views of the same mouse (from below and from the side), and two epochs in the same behavioral session. The first epoch is prior to the injection of ethanol (1.5 g/kg IP), during which the mouse displays normal locomotor behavior. The second epoch is approximately 10-15 minutes after the injection of ethanol, during which it shows prominent dystonic movements involving the trunk, limbs, and tail.

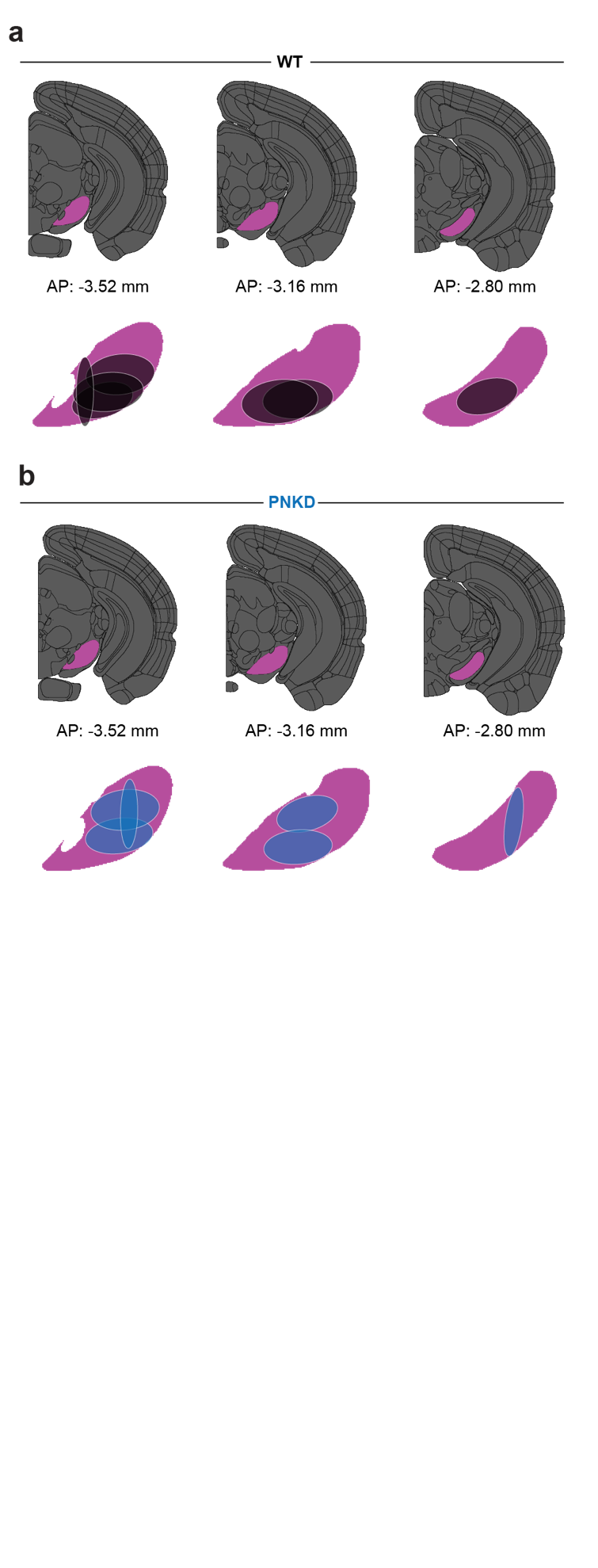
**Fig. S1. Distribution of electrode recording channels in the SNr**

**(Related to Fig. 1)**

**a,b,** Location of electrode array channels and high-density probe channels within the SNr of wild-type (WT) (a, N = 7 mice, n = 75 neurons) and paroxysmal nonkinesigenic dyskinesia (PNKD) mice (b, N = 6, n = 46). Location of electrode channels and probe channels were confirmed by electrolytic lesion or DiI lipophilic dye, respectively. These locations were registered to the Allen Mouse Brain Atlas and only single-unit recordings from channels located within the SNr were retained for analysis.

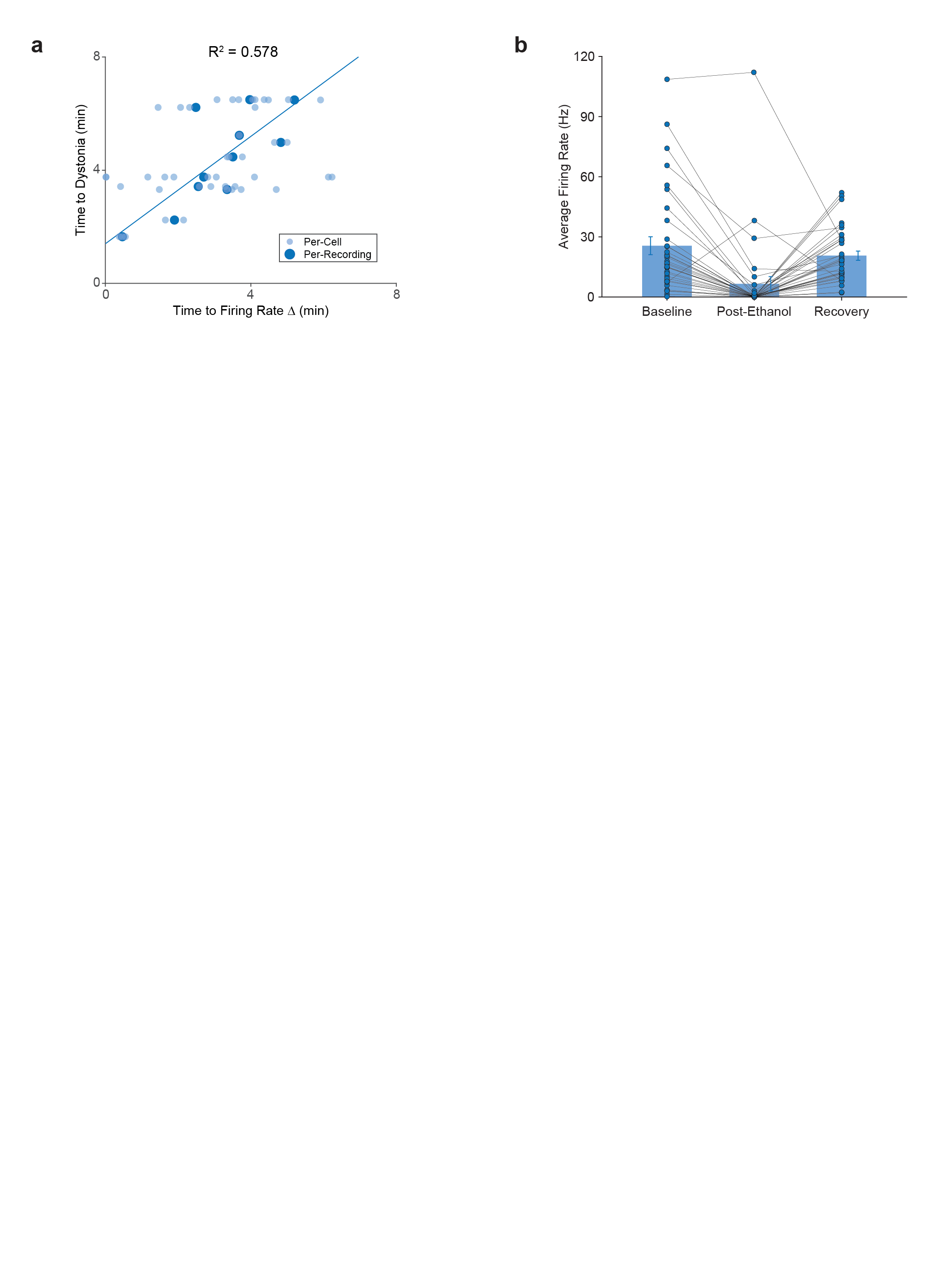
**Fig. S2. SNr Neuron Activity During the Onset and Offset of Dystonia**

**(Related to Fig. 1)**

**a,** Correlation between the time at which the SNr firing rate fell below the 99% confidence interval and the onset of dystonic movement. Small, light blue circles represent values for individual cells across all recordings (N = 6, n = 44; Pearson correlation coefficient; *r = 0.471; R^2^ = 0.222; P = 0.0013*). Larger, dark blue circles and the line of best fit correspond to the correlation between an average of the firing rate change for an individual recording as a function of dystonia onset (N = 6, n = 11; Pearson correlation coefficient; *r = 0.760; R^2^ = 0.578; P = 0.0066*). **b,** In recordings dystonia was found to resolve completely, comparison of PNKD SNr neuron firing rates during baseline (30 minutes prior to ethanol injection), post-ethanol (10–70 minutes after ethanol injection), and recovery (ends of recording during which dystonia had resolved for at least 10 minutes) periods (N = 5, n = 34), repeated measures ANOVA, *F*(2,66) = 12.377, *P* < 0.0001, post-hoc Wilcoxon signed-rank test: baseline vs post-ethanol, *P* < 0.0001 *;* baseline vs recovery, *P* = 0.7518; post-ethanol vs recovery, *P* < 0.0001). Data shown as mean ± SEM.

**Table S2**. Excitability and Action Potential Shape Parameters*^b^*

| **Parameter** | **N (animals)** | **n (cells)** | **Mean ± SEM** | **Statistical Test** | ***P*** |
| --- | --- | --- | --- | --- | --- |
| Input Resistance (MOhm) | WT: 6 PNKD: 7 | WT: 71  PNKD: 101 | WT: 161.21 ± 6.729  PNKD: 186.79 ± 8.056 | MWU | 0.0612 |
| Membrane Potential (mV) | WT: 6 PNKD: 7 | WT: 71 PNKD: 100 | WT: -48.00 ± 0.799  PNKD: -50.04 ± 0.793 | MWU | *0.0138 |
| First Action Potential Width (ms) | WT: 6 PNKD: 7 | WT: 69 PNKD: 96 | WT: 0.6528 ± 0.0161  PNKD: 0.6079 ± 0.0174 | MWU | *0.0209 |
| First Action Potential Threshold (mV) | WT: 6 PNKD: 7 | WT: 69 PNKD: 96 | WT: -43.76 ± 0.6993  PNKD: -42.77 ± 0.6182 | MWU | 0.6366 |
| First Action Potential AHP Amplitude (mV) | WT: 6 PNKD: 7 | WT: 69 PNKD: 96 | WT: -65.43 ± 0.7823  PNKD: -65.96 ± 0.5618 | MWU | 0.3977 |
| Average Action Potential Width (ms) | WT: 6 PNKD: 7 | WT: 66 PNKD: 79 | WT: 0.7221 ± 0.0195  PNKD: 0.6932 ± 0.0214 | MWU | 0.2319 |
| Average Action Potential Threshold (mV) | WT: 6 PNKD: 7 | WT: 66 PNKD: 79 | WT: -32.7052 0.7780  PNKD: -32.3807 ± 0.5858 | MWU | 0.8629 |
| Average Action Potential AHP Amplitude (mV) | WT: 6 PNKD: 7 | WT: 66 PNKD: 79 | WT: -58.9580 ± 0.9609  PNKD: -58.7251 ± 0.6430 | MWU | 0.7613 |

*^b^*MWU: Mann-Whitney *U* test; AHP: Afterhyperpolarization.

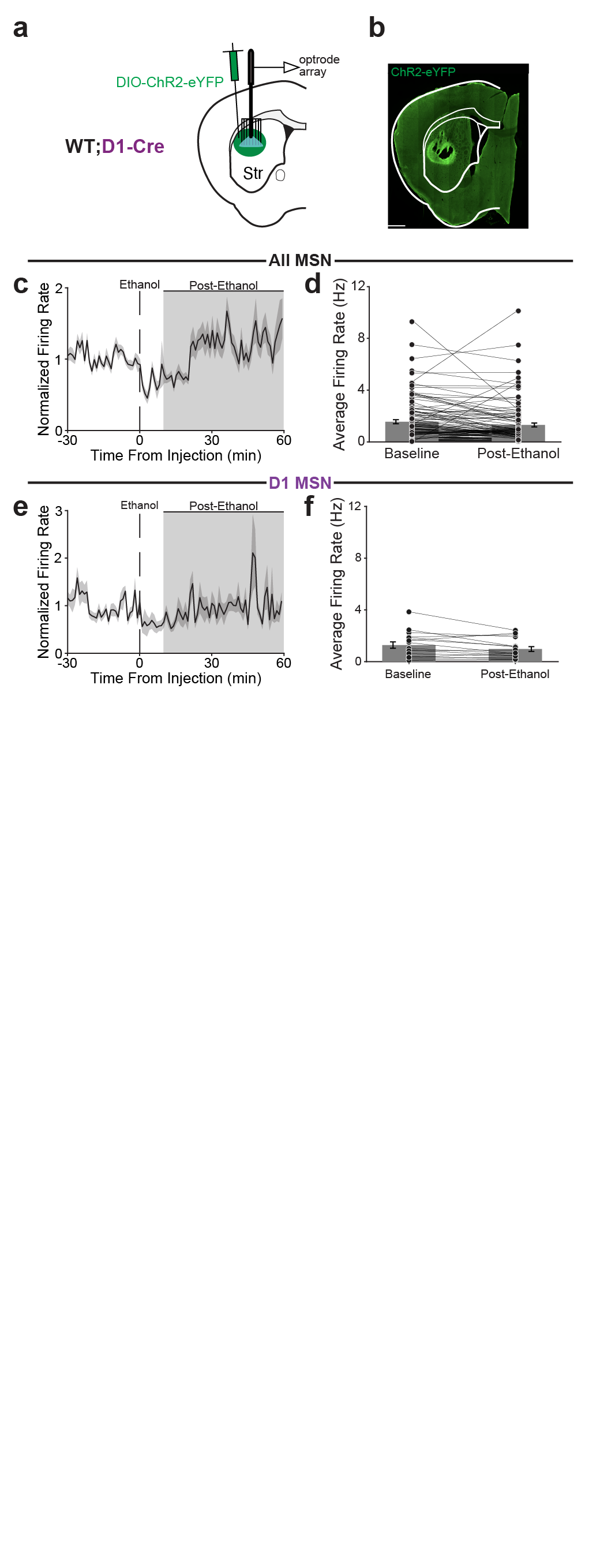
**Fig. S3: MSN activity is modestly reduced after ethanol administration in healthy mice**

**(Related to Fig. 3)**

**a,** Schematic showing optrode recording configuration and injection of AAV encoding DIO-ChR2-eYFP in the DLS of WT;D1-Cre or WT;A2a-Cre mice. **b,** Representative histological confirmation of ChR2-eYFP viral expression and location of electrode confirmed by electrolytic lesion. White scale bar represents 1 mm. **c,** Average normalized firing rate of all MSNs before and after ethanol injection in WT mice. **d,** Comparison of MSN average firing rates during baseline and post-ethanol periods (N = 10, n = 139, Wilcoxon signed-rank test, *P* = 0.0002). **e,** Average normalized firing rate of D1-MSNs before and after ethanol injection in WT mice. **f,** Comparison of D1-MSN average firing rates during baseline and post-ethanol periods (N = 2, n = 17, Wilcoxon signed-rank test, *P* = 0.0615). All data shown as mean ± SEM.

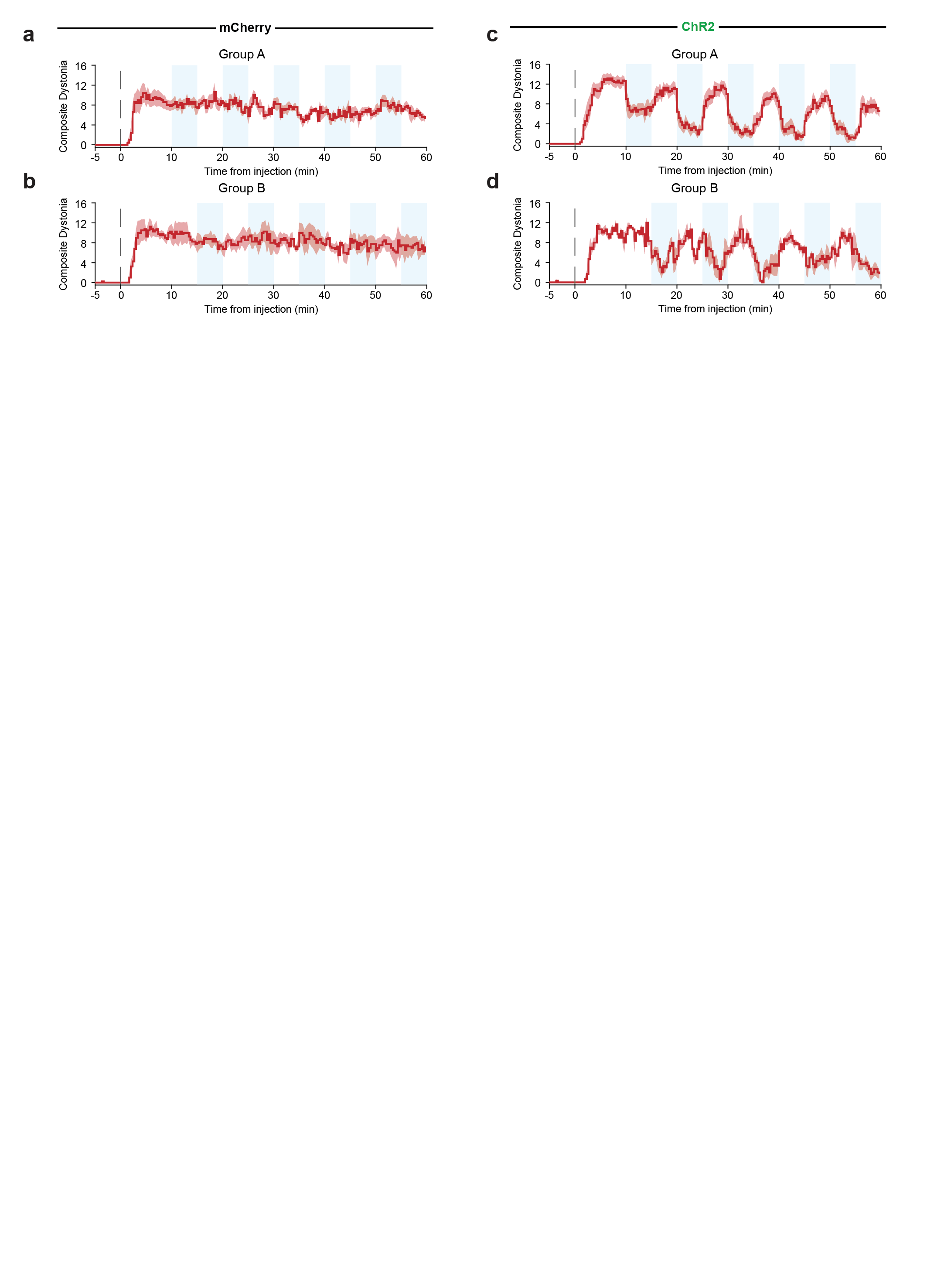
**Fig. S4: Time course of PNKD dystonia during optogenetic rescue across all experimental groups (Related to Fig. 8)**

**a–d,** Dystonia severity in mice expressing mCherry (a,b) or ChR2-eYFP (c,d) in the dorsal striatum. Blue light was delivered bilaterally into the SNr at 4 Hz, 8 ms pulses, and 3–6 mW in alternating 5-minute light-off and light-on epochs (five each). Mice were assigned to one of two groups differing in which epoch came first: Group A received a light-on epoch first (beginning at 10 minutes after ethanol injection) and Group B received a light-off epoch first (followed by light-on epoch at 15 minutes). **a,** Average dystonia severity time course for mCherry-expressing, Group A mice (N = 5). **b,** Average dystonia severity time course for mCherry-expressing, Group B mice (N = 4). **c,** Average dystonia severity time course for ChR2-expressing, Group A mice (N = 7). **d,** Average dystonia severity time course for ChR2-expressing, Group B mice (N = 3). All data shown as mean ± SEM.

**Supplementary Video 2. Behavior of PNKD mouse during light-off and light-on epochs in a dystonic attack.** This PNKD mouse was injected with bilateral ChR2 in the dorsal striatum, and optical fibers were placed in the bilateral SNr. Ethanol (1.5 g/kg) was injected IP approximately 30 minutes prior to the epochs captured here. In the first epoch, no light was delivered to the SNr. In the second epoch, blue (473 nm) light was delivered to the bilateral SNr in 8 msec pulses, delivered at 4 Hz, 3 mW.
